# Liver Lymphatic Dysfunction as a Driver of Fibrosis and Cirrhosis Progression

**DOI:** 10.1101/2025.01.11.632552

**Authors:** Jain Jeong, Shao-Jung Hsu, Daiki Horikami, Teruo Utsumi, Yilin Yang, Nikolai Arefyev, Xuchen Zhang, Shi-Ying Cai, James L. Boyer, Rolando Garcia-Milan, Masatake Tanaka, Matthew J. McConnell, Hui-Chun Huang, Yasuko Iwakiri

## Abstract

The liver lymphatic system plays a critical role in maintaining interstitial fluid balance and immune regulation. Efficient lymphatic drainage is essential for liver homeostasis, but its role in liver disease progression remains poorly understood. In cirrhosis, lymphangiogenesis initially compensates for increased lymph production, but impaired lymphatic drainage in advanced stages may lead to complications such as ascites and portal hypertension. This study aimed to evaluate how liver lymphatic dysfunction affects disease progression and to assess therapeutic strategies. Using a surgical model to block liver lymphatic outflow, we found that impaired drainage accelerates liver injury, fibrosis, and immune cell infiltration, even in healthy livers. Mechanistically, enhanced TGF-β signaling in liver lymphatic endothelial cells (LyECs) contributed to reduced lymphatic vessel (LV) density and function in late-stage decompensated cirrhosis. This dysfunction was linked to the progression from compensated to decompensated cirrhosis, particularly in patients with primary sclerosing cholangitis (PSC). Conversely, liver-specific overexpression of VEGF-C via AAV8 improved lymphatic drainage, restored LV density, reduced fibrosis, mitigated liver injury, and alleviated portal hypertension in cirrhotic rats. These findings establish impaired liver lymphatic function as a pivotal driver of cirrhosis progression and identify VEGF-C as a promising therapeutic target to prevent decompensation.

## INTRODUCTION

The liver lymphatic system plays a critical role in maintaining interstitial fluid (lymph) balance(1, 2) and facilitating immune surveillance(3–6). As one of the largest lymph-producing organs, the liver contributes 25–50% of the lymph flow through the thoracic duct(7, 8). Efficient lymphatic drainage is critical for liver homeostasis, yet the impact of impaired liver lymphatic function on liver health and disease progression remains incompletely understood.

In patients with cirrhosis, lymph production increases by up to 6-fold(9, 10), accompanied by lymphangiogenesis, the formation of new lymphatic vessels (LVs) in the liver and gut, to manage the heightened lymphatic load(3, 4). However, as cirrhosis progresses, lymphatic drainage capacity becomes overwhelmed, leading to lymph leakage into the peritoneal cavity (i.e., ascites)(11–14). A study in rats with carbon tetrachloride (CCl_4_)-induced liver fibrosis and cirrhosis showed that while functional LV density increases in early fibrosis, it tends to decrease in late-stage fibrosis/cirrhosis(1), suggesting an impairment of lymphatic drainage in advanced disease stages.

Cirrhosis is closely linked to portal hypertension, a condition that evolves from a compensated stage, which is largely asymptomatic, to a decompensated stage characterized by severe complications such as ascites(15, 16). Portal hypertension arises primarily from increased intrahepatic resistance due to fibrosis and may be exacerbated by impaired liver lymphatic drainage. For instance, in anesthetized cats, blocking liver lymphatic drainage through LV ligation near liver-draining lymph nodes significantly increased hepatic venous pressure and sinusoidal lymph production(17). This finding suggests that impaired liver lymphatic drainage alone is sufficient to elevate intrahepatic resistance and hydrostatic pressure, worsening portal hypertension. However, the broader implications of impaired liver lymphatic drainage on liver homeostasis, function and disease progression remain to be fully elucidated. Thus, this study aims to deepen our understanding of the role of the liver lymphatic system in maintaining homeostasis, supporting function, and influencing disease progression.

Lymphangiogenic factors such as vascular endothelial growth factor (VEGF)-C and VEGF-D promote lymphatic drainage by increasing LV numbers and area through activation of VEGF receptor 3 (VEGFR3, also known as Flt4)(18–22). Recent evidence highlights the therapeutic potential of VEGF-C/D-driven lymphangiogenesis in liver diseases, including MAFLD, MASH, and ischemic-reperfusion injury(20, 23–28). Notably, increased mesenteric lymphatic drainage with recombinant VEGF-C (Cys156Ser) nanoparticle has been shown to reduce portal pressure and ascites in cirrhotic rats(29). However, this approach, which targets mesenteric rather than liver lymphatics, did not reduce liver fibrosis(29), leaving its effect on fibrosis and cirrhosis unclear.

In this study, we used a surgical model to block liver lymphatic outflow and demonstrated that impaired drainage accelerates liver injury, fibrosis, and immune cell infiltration, even in normal liver. Additionally, we observed decreased LVs and reduced lymphatic drainage function in compensated cirrhosis, driven by enhanced TGF-β signaling in liver lymphatic endothelial cells (LyECs). This impairment promotes progression from the compensated to the decompensated stage in both mice and patients with cirrhosis, particularly in those with primary sclerosing cholangitis (PSC). Conversely, enhanced lymphatic drainage with liver-targeted AAV8-VEGF-C significantly reduced fibrosis, mitigated injury, and alleviated portal hypertension in cirrhotic rats. These findings establish impaired liver lymphatic function as a key driver of disease progression from compensated to decompensated cirrhosis and identify VEGF-C as a promising therapeutic candidate.

## RESULTS

### Blocking liver lymphatic drainage leads to increased liver injury and fibrosis

We surgically blocked liver lymphatic drainage and evaluated its impact on the liver (**Figure 1A**), referring to this surgical procedure as lymphatic obstruction (LO) surgery afterwards. Three days post-LO surgery, we observed a significant increase in liver weight (**Figure 1B**), indicative of fluid (lymph) retention due to impaired lymphatic drainage. Additionally, spleen weight increased by 1.2-fold (p<0.01), while body weight remained unchanged compared to sham-operated mice (sham vs. LO: 24.0 ± 0.13g vs. 23.8 ± 0.10g, p=0.33). LO surgery resulted in a significant increase in lymphatic vessel (LV) number and area (**Figure 1C, D**), as well as mild elevations in liver plasma ALT levels (1.8-fold, p<0.05), a marker of liver injury (**Figure 1E**), liver fibrosis (1.4-fold, p<0.05) (**Figure 1F, G**), and hepatic stellate cell (HSC) activation (**Figure 1H-K**).

**Figure 1.**
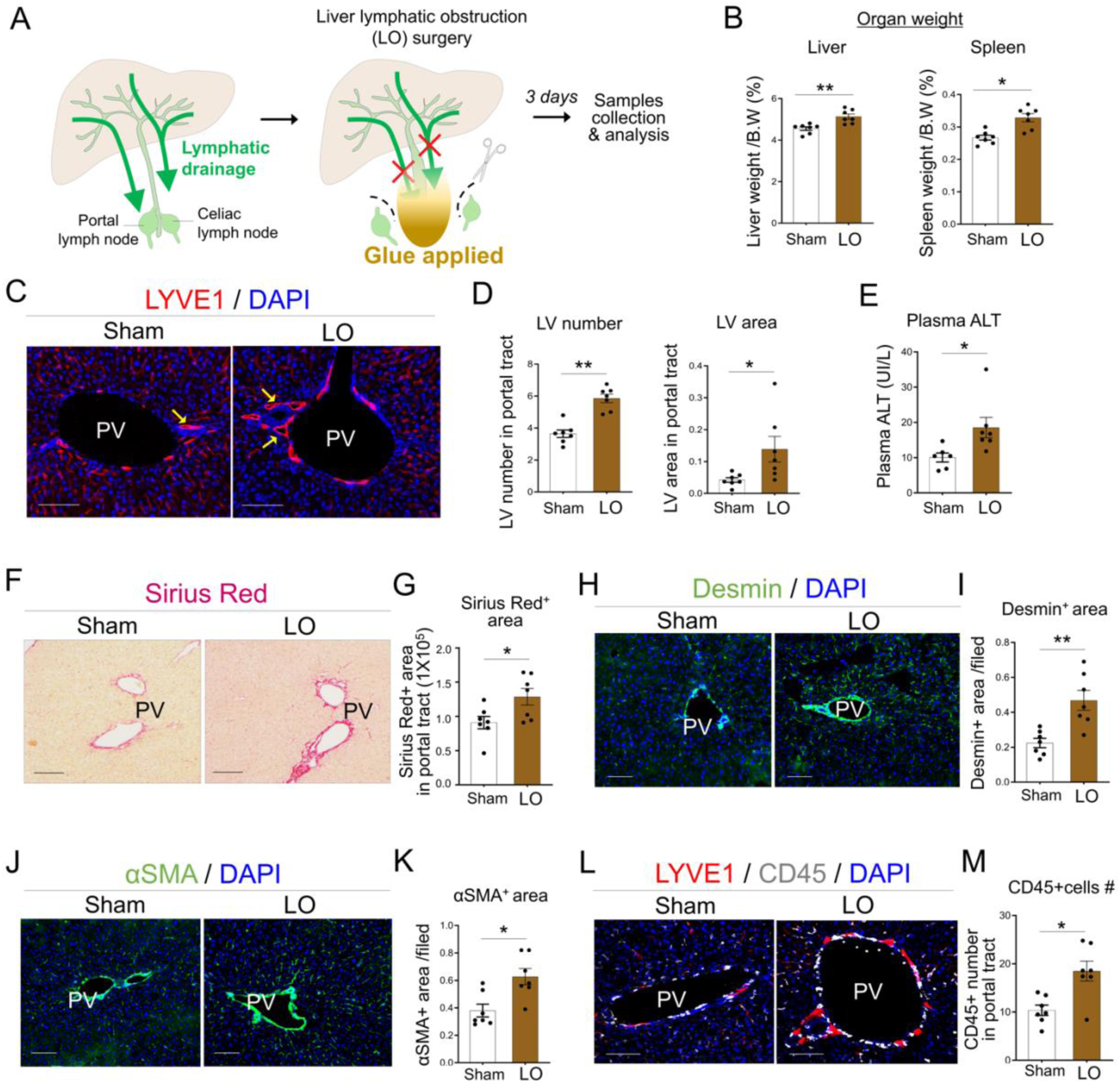
Blocking the liver lymphatic drainage leads to increased liver injury and fibrosis. (A) Liver lymphatic obstruction (LO) surgery. Three days after the lymph blockage, mice were sacrificed for analysis. (B) Liver and spleen weights in mice 3 days after liver LO surgery (n=7) or sham surgery (n=7). B.W.: body weight. (C) Immunofluorescence (IF) images of LYVE-1 [lymphatic vessels (LVs); red], and DAPI in mice 3 days after LO (n=7) or sham surgery (n=7). Arrows indicate LVs. (D) Quantification of LV number and area in mice 3 days after LO (n=7) or sham surgery (n=7). (E) Liver damage evaluated by ALT value in plasma of mice 3 days after LO (n=7) or sham surgery (n=7). (F-K) Evaluation of liver fibrosis. Representative Sirius red staining images (F) and its quantification (G) in livers from mice 3 days after LO (n=7) or sham surgery (n=7). (H) IF images of Desmin [a marker of hepatic stellate cells (HSCs); green], and (I) its quantification in mice 3 days after LO (n=7) or sham surgery (n=7). (J) IF images of α-SMA (a marker of activated HSCs and smooth muscle cells; green), and (K) its quantification in mice 3 days after LO (n=7) or sham surgery (n=7). (L) Representative IF images of CD45 (a marker of lymphocytes; white), LYVE1 and DAPI and (M) quantification in mice 3 days after LO (n=7) or sham surgery (n=7). *P < 0.05, and **P < 0.01. Scale bars: 100 μm. PV, portal vein.

Lymphatic vessels (LVs) facilitate immune cell egress(30). Consistent with this, CD45-positive leukocytes were significantly increased (1.8-fold, p<0.05) in peri-portal areas of livers from LO mice (**Figure 1L, M**). These findings suggest that effective liver lymphatic drainage is crucial in mitigating liver injury, fibrosis, and immune cell infiltration.

### Inhibition of VEGF-C/VEGFR3 signaling by MAZ51 aggravates liver fibrosis in cholestatic mice

To examine the role of lymphatic system in liver fibrosis, we used bile duct ligation (BDL) mouse model, a well-established experimental model of cholestatic liver disease. We chose this model, because it develops biliary fibrosis and inflammation primarily in the portal tract area(31), where most of intrahepatic LVs are located. Liver fibrosis, measured by Sirius red staining, increased significantly one week post-BDL (BDL1w; 2.0-fold, p<0.001), but showed no change in hydroxyproline level (p=0.167) (**Figure 2A-C**). Fibrosis progression was time-dependent, with Sirius red-positive areas and hydroxyproline levels increasing markedly at 2 weeks (BDL2w) and peaking at 4 weeks (BDL4w) (Sirius red: 4.4-fold, p<0.001; hydroxyproline: 2.0-fold, p<0.01) (**Figure 2B, C**). Serum ALT levels followed a similar trend (**Figure 2D**). As fibrosis progressed, hallmarks of decompensated cirrhosis (DC) including severe portal hypertension, portosystemic shunt and hepatopulmonary syndrome, emerge at 4-5 weeks post-BDL surgery(32, 33).

**Figure 2.**
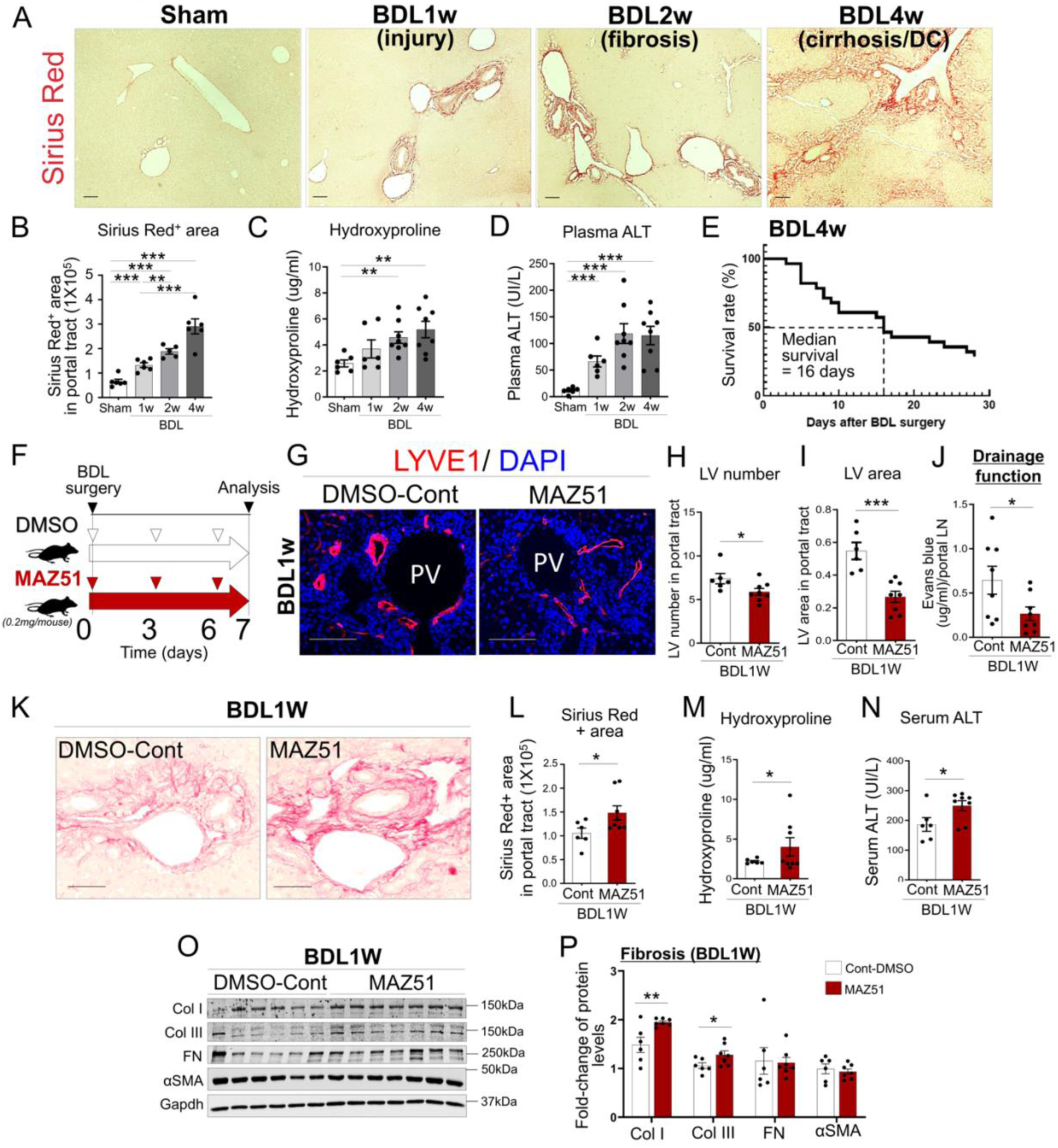
Inhibition of VEGF-C/VEGFR3 signaling by MAZ51 decreases liver lymphatic drainage, leading to increased liver injury and fibrosis in cholestatic mice. (A-D) Pathological stages of the livers in mice with bile duct ligation (BDL). Representative Sirius Red staining images (A) and its quantification (B), hydroxyproline level (C) and serum ALT levels (D) in sham (n=6), 1-week(n=6), 2-week(n=5), and 4-week(n=7) BDL mice. (E) Kaplan-Meier survival curve showing a statistically significantly decreased in median survival 16 days after BDL surgery (n=28). (F) Experimental scheme of MAZ51 injection in mice. DMSO(control) or MAZ51 was injected subcutaneously (0.2mg/mouse). Arrow heads indicate the day of injection. (G) Immunofluorescence (IF) images of LYVE-1 [lymphatic vessel (LV); red] and DAPI in livers from 1-week BDL mice given DMSO (control) or MAZ51. (H, I) Quantification of LV numbers and area in livers from 1-week BDL mice treated with control-DMSO(n=6) and MAZ51(n=8). (J) Liver lymphatic drainage test. Evans blue (EVB) dye injected directly into the liver was measured in liver draining portal lymph nodes (pLNs) by microplate reader (650nm) as an indicator of liver lymphatic drainage function, 7 days after the injection of DMSO control (n=6) or MAZ51 (n=8). (K-N) Evaluation of liver fibrosis and damage. Representative Sirius red staining images (K) and its quantification (L), as well as hydroxyproline levels (M) and serum ALT levels (N) in livers from 1-week BDL mice treated with control-DMSO(n=6) or MAZ5(n=8). (O&P) Fibrotic maker expression including collagen I (ColI), collagen III (Col III), fibronectin 1 (FN1) and α SMA as indicated by western blotting. n=6 (control-DMSO) and n=8 (MAZ5). *P < 0.05, **P < 0.01, and ***P < 0.001. Scale bars: 100 μm. PV, portal vein.

Survival rates declined sharply, with approximately 50% of mice surviving by day 16 and only 30% surviving by 4 weeks (**Figure 2E**). Based on these findings, we defined the 4-week (28-day) time point post-BDL surgery (BDL4w) as the late-stage DC stage(34). In contrast, the 2-week time point (BDL2w) represented an intermediate stage between advanced fibrosis and compensated cirrhosis, while the 1-week time point (BDL1w) reflected the early onset of fibrosis associated with cholestatic liver injury.

To assess whether impaired lymphangiogenesis accelerates fibrosis, we administered the VEGFR3 inhibitor MAZ51 (0.2mg/mouse) or DMSO (control) to BDL1w mice (**Figure 2F**). MAZ51 treatment reduced LV number and area (**Figure 2H, I**) and decreased liver lymphatic drainage (∼42.2 %, p<0.05) (**Figure 2J**). This impairment corresponded with increased Sirius red-positive fibrotic area (1.3-fold, p<0.05) and hydroxyproline levels (1.8-fold, p<0.05), and ALT levels (1.3-fold, p<0.05) (**Figure 2K-N, Supplementary Figure 1A**). Fibrosis markers, such as collagen I (CoI I) and III (Col III), were also elevated (**Figure 2O, P**). MPO-positive immune cell infiltration increased (**Supplementary Figure 1B**). Body, liver, and spleen weights remained unchanged between groups (**Supplementary Figure 2A&B**). These findings indicate that impaired liver lymphatic drainage exacerbates liver injury and fibrosis.

### VEGF-C overexpression enhances liver lymphangiogenesis and lymphatic drainage, preventing injury, fibrosis and LSEC capillarization

To determine whether enhancing lymphangiogenesis and liver lymphatic drainage prevents liver disease progression, we overexpressed VEGF-C using AAV8-VEGF-C and AAV8-GFP as control (**Figure 3 A**). VEGF-C mRNA (3.0-fold, p<0.001) and protein expression (1.2-fold, p<0.05) increased significantly in VEGF-C-treated BDL2w mice compared to control (**Figure 3B, C**). Interestingly, VEGF-C treatment decreased spleen weight ratio in BDL2w mice (p<0.05) compared to control mice (**Figure 3D**), while there were no differences in body and liver weights between groups (**Figure 3D**). VEGF-C treatment increased LV number and area (**Figure 3E-G**) without affecting angiogenesis (**Supplementary Figure 3B**) and bile duct area (**Supplementary Figure 3C**), and enhanced liver lymphatic drainage function (∼224%, p<0.05)(**Figure 3H**). It reduced fibrosis (Sirius red area, −1.4-fold, p<0.05; hydroxyproline level, −1.4-fold, p<0.05) and liver injury as indicated by ALT levels (−1.5-fold, p=0.051)(**Figure 3I-L, Supplementary Figure 3A**). Fibrosis markers, such as collagen I (CoI I) and III (Col III) expressions, were decreased by VEGF-C treatment (**Figure 3M, N**).

**Figure 3.**
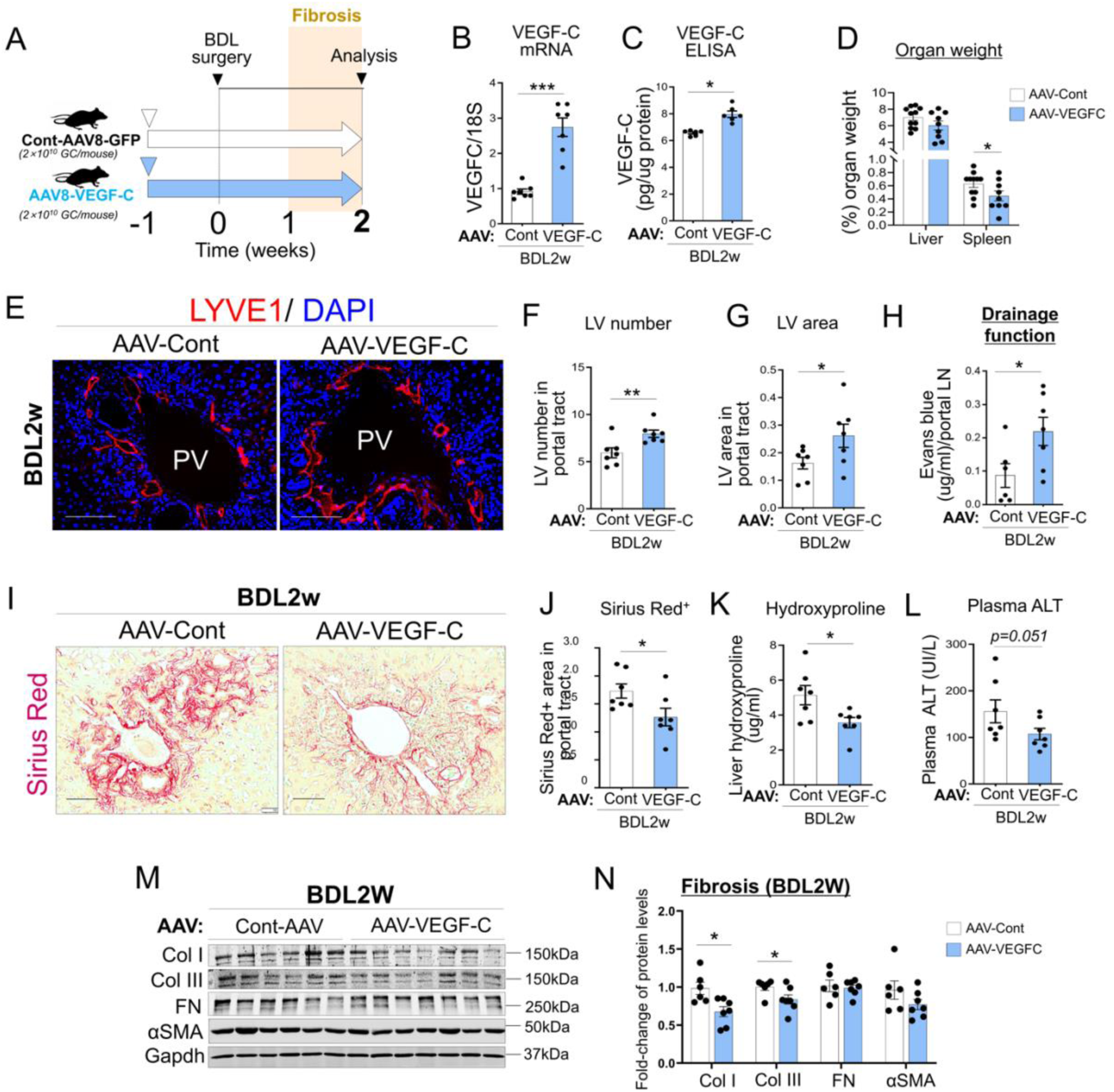
AAV8-medicated VEGF-C overexpression increases liver lymphatic drainage function, leading to decreased liver injury and fibrosis in cholestatic mice. (A) Experimental scheme of AAV8 injection in cholestatic mice. Retro-orbital injection of AAV8-VEGF-C or AAV8-GFP(control) (1.5×10^11^ GC/mouse) was conducted 7 days before BDL surgery. Two weeks after BDL surgery, liver samples were collected for analysis. (B, C) VEGF-C mRNA and protein levels in liver lysates from 2-week BDL mice treated with AAV8-GFP (n=7) or AAV8-VEGF-C (n=7). (D) Liver and spleen weights in mice treated with AAV8-GFP (Control, n=11) or AAV8-VEGF-C (n=9). (E) Immunofluorescence (IF) images of LYVE-1 [Lymphatic vessel (LV); red) and DAPI as well as its quantification (F, G) of LV number and area in livers from 2-week BDL mice treated with AAV8-GFP(n=7) or AAV8-VEGF-C(n=7). (H) Liver lymphatic drainage function in 2-week BDL mice given AAV8-GFP (n=6) or AAV8-VEGF-C (n=7). (I-L) Evaluation of liver fibrosis and damage. Representative Sirius Red staining images (I) and its quantification (J) as well as hydroxyproline level (K), serum ALT levels (L) in livers from 2-week BDL mice treated with AAV8-GFP(n=7) or AAV8-VEGF-C(n=7). (M) Expression of liver fibrotic makers: collagen I (ColI), collagen III (Col III), fibronectin 1 (FN1), and α SMA by western blotting and the quantification (N). AAV8-GFP (n=7) and AAV8-VEGF-C (n=7). *P < 0.05, **P < 0.01, and ***P < 0.001. Scale bars: 100 μm. PV, portal vein.

VEGF-C treatment also decreased CD68-positive macrophage infiltration (**Supplementary Figure 4**) and restored liver sinusoidal endothelial cell (LSEC) functionality by reducing CD34 expression (**Supplementary Figure 5B, C**). VEGF-C-driven recovery of LSEC function can be explained by its expression of VEGFR3. Re-analysis of single-cell RNA sequencing data(35) indicated that while VEGFR-3 expression was the highest in liver lymphatic endothelial cells (LyECs) in mice, a considerable expression was observed in LSECs (**Supplementary Figure 5A**).

### VEGF-C overexpression ameliorates portal hypertension in cholestatic rats

To determine whether enhanced liver lymphatic drainage via AAV8-VEGF-C could mitigate the progression of late-stage liver cirrhosis, we evaluated hemodynamic parameters, including intrahepatic resistance, portal pressure in a late-stage cirrhotic BDL rat model (rBDL4w), characterized by decompensated cirrhosis(18) (**Figure 4A**). Rats were chosen for this study due to the higher consistency and reproducibility of hepatic hemodynamic measurements compared to mice(36, 37). VEGF-C overexpression in the liver significantly reduced fibrosis progression in late-stage cirrhotic rats (**Figure 4B**), and decreased hepatic vascular resistance and portal pressure by 20% (p<0.05 and p<0.01, respectively) (**Figure 4C, D**), without affecting systemic circulation (**Figure 4E, F**), cardiac function (**Figure 4G**), superior mesenteric artery flow, or collateral circulation (**Figure 4H-J**). Collectively, these findings demonstrate that VEGF-C-driven enhancement of liver lymphatic drainage ameliorates portal hypertension in late-stage cirrhotic rata (**Figure 4K**).

**Figure 4.**
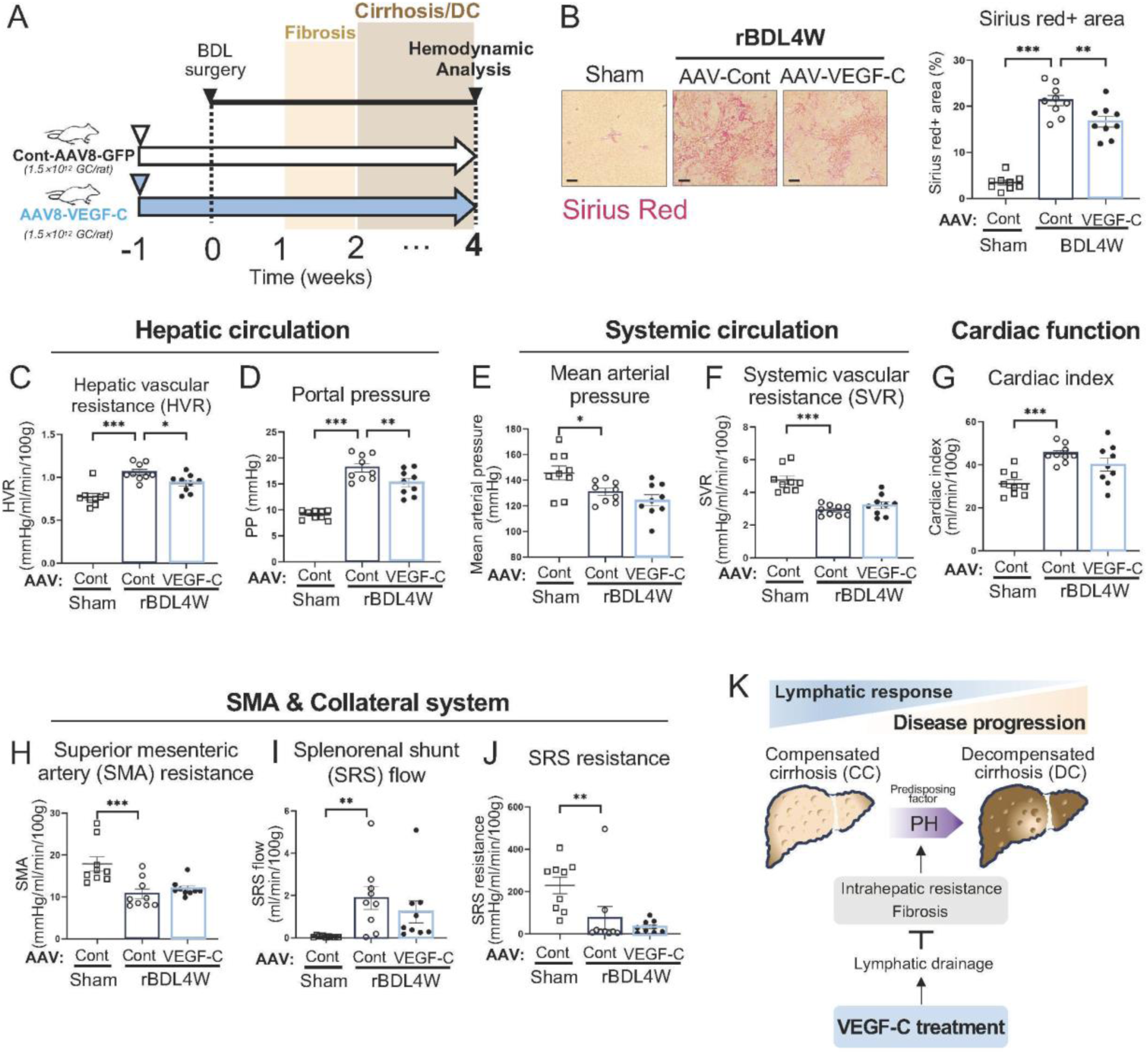
AAV8-mediated VEGF-C overexpression ameliorates portal hypertension in cholestatic rats. (A) Experimental scheme of AAV8 injection in rats. Retro-orbital injection was performed for AAV8-VEGF-C or AAV8-GFP(control) (1.5×10^11^ GC/rat) 7 days before BDL surgery. Four weeks after BDL surgery, liver samples were collected for analysis. Evaluation of liver fibrosis (B) and hemodynamic characterization performed for Hepatic vascular resistance (C); portal pressure (D); mean arterial pressure (E); systemic vascular resistance (SVR)(F); cardiac index(G); superior mesenteric artery (SMA) resistance(H); splenorenal shunt (SRS) flow(I); and SRS resistance(J), measured 4 weeks after BDL or sham surgery. Sham-operated AAV8-GFP rat (n=9), 4-week BDL rats treated with AAV8-VEGF-C (n=9) or AAV8-GFP(n=9). *P < 0.05, **P < 0.01, and ***P < 0.001. PH, portal hypertension.

### Liver lymphatic vessels decease during progressing to decompensated cirrhosis in mice and human patients

To test our hypothesis that increased lymphangiogenesis protects against fibrosis, while impaired lymphangiogenesis and lymphatic drainage facilitate disease progression, we evaluated lymphangiogenesis across different stages of fibrosis and cirrhosis in mice (**Figure 5A-E**). The number and area of LVs peaked in BDL2w mice (fibrosis/cirrhosis stage) with significant increases (2.9-fold, p<0.001 and 4.1-fold, p<0.01 respectively), but declined in BDL4w mice (late-stage cirrhosis) by −1.4-fold (p<0.05) and −1.9-fold (p<0.05), respectively (**Figure 5A-C**). Similarly, the percentage of proliferating cell nuclear antigen (PCNA)-positive LyECs, indicative of active lymphangiogenesis, was highest at 2 weeks post-BDL surgery, but decreased by 4 weeks (**Supplementary Figure 6A, B**). VEGF-C, a key lymphangiogenic factor, showed elevated protein levels in BDL2w and BDL4w mice compared to sham mice (**Figure 5D**), and its mRNA levels positively correlated with LV numbers in BDL livers (**Figure 5E**).

**Figure 5.**
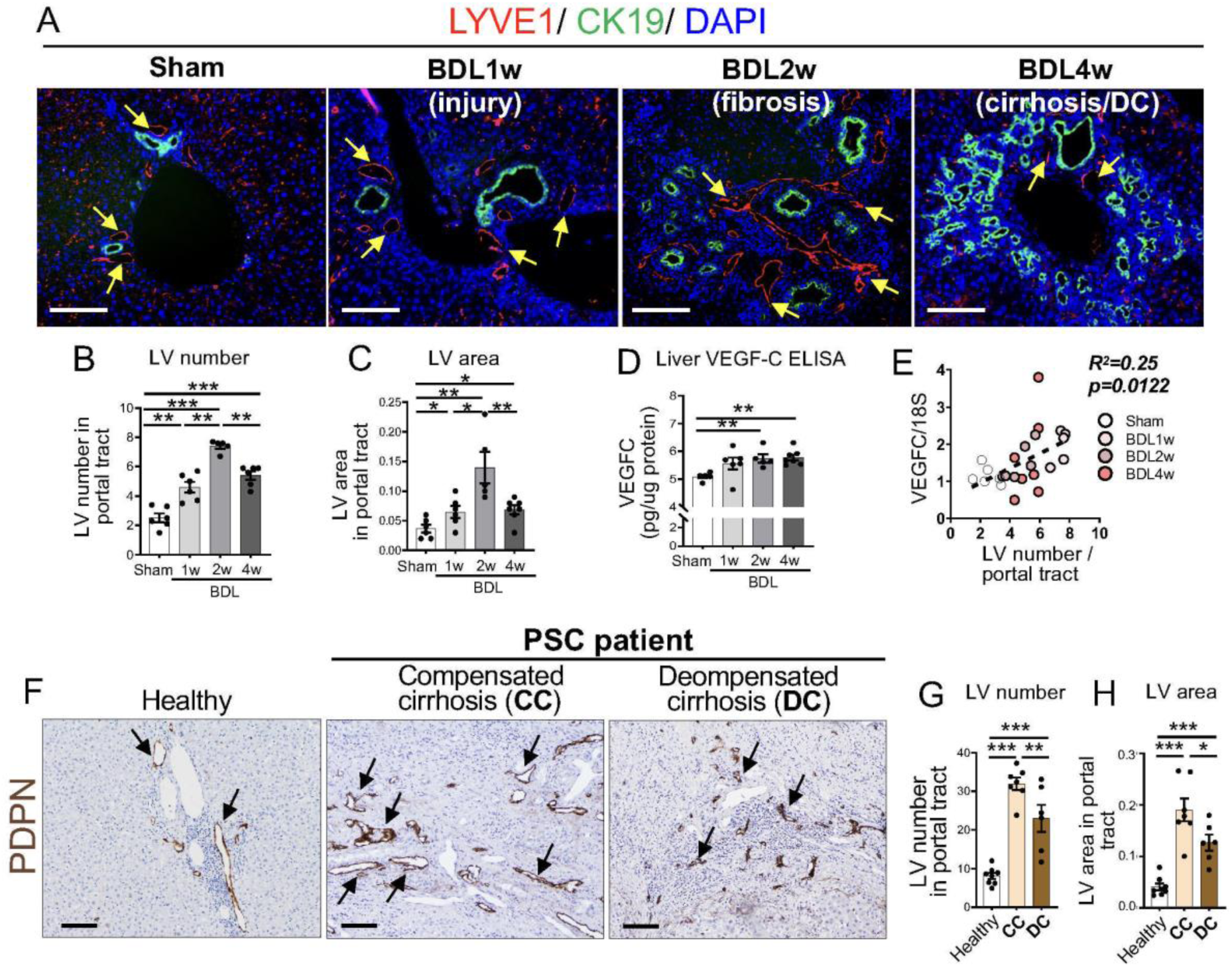
Liver lymphatic vessel (LV) numbers and area increased in fibrosis/cirrhosis but decreased in decompensated cirrhosis in cholestatic mice and patients with primary sclerosing cholangitis (PSC) (A) Representative immunofluorescence (IF) images in livers from sham, 1-week, 2-week, 4-week BDL mice. LYVE-1 [Lymphatic vessels (LVs)]; red), CK19 (bile duct; green), and DAPI. Arrows indicate LVs. (B, C) Quantification of LV number and area in sham (n=6), 1-week(n=6), 2-week(n=5), and 4-week(n=7) BDL mouse livers. (D) VEGF-C protein levels in liver lysates from sham(n=6), 1-week(n=6), 2-week(n=5), and 4-week(n=7) BDL mice. (E) Positive correlation between liver VEGF-C mRNA expression and LV numbers (n=24). (F) Representative LV images [podoplanin (PDPN); a LV marker) in livers from healthy controls and PSC patients classified into compensated cirrhosis (CC) and decompensated cirrhosis (DC). Arrows indicate LVs. (G, H) Quantification of LV number and area in livers from healthy controls (n=8) and PSC patients who are CC(n=7) or DC(n=6). *P < 0.05, **P < 0.01, and ***P < 0.001. Scale bars: 100μm.

In parallel, we analyzed liver specimens from 13 patients with primary sclerosing cholangitis (PSC), dividing them into compensated cirrhosis (CC) and late-stage decompensated cirrhosis (DC) groups based on clinical history of complications or medications indicating ascites or hepatic encephalopathy(38)(**Supplementary Table 1**). Podoplanin (PDPN) a marker for LVs, revealed that LV number and area were highest in compensated cirrhosis, but declined in late-stage decompensated cirrhosis (**Figure 5F-H and Supplementary Figure 7**). Collectively, these findings suggest that the regression of LVs during late-stage cirrhosis contributes to the transition from compensated to decompensated cirrhosis.

### Transcriptomic analysis reveals increased TGF-β signaling in liver lymphatic endothelial cells in late-stage cirrhosis

To gain insight into the molecular mechanisms underlying changes in LyEC function during fibrosis and late-stage decompensated cirrhosis, we performed RNA sequencing on liver LyECs isolated from livers of mice subjected to sham (n=3), 2-week (BDL2w, n=3), or 4-week (BDL4w, n=3) BDL surgery (**Figure 6A**). Unsupervised clustering of transcriptomic data revealed distinct gene expression patterns in LyECs at different stages of fibrosis and cirrhosis (**Figure 6B**). In total, 207 genes were differentially expressed in LyECs from BDL2w mice compared to sham, while 554 genes were differentially expressed in LyECs from BDL4w mice compared to sham (**Figure 6B**). Of these, 74 genes were altered in LyECs from both BDL2w and BDL4w groups.

**Figure 6.**
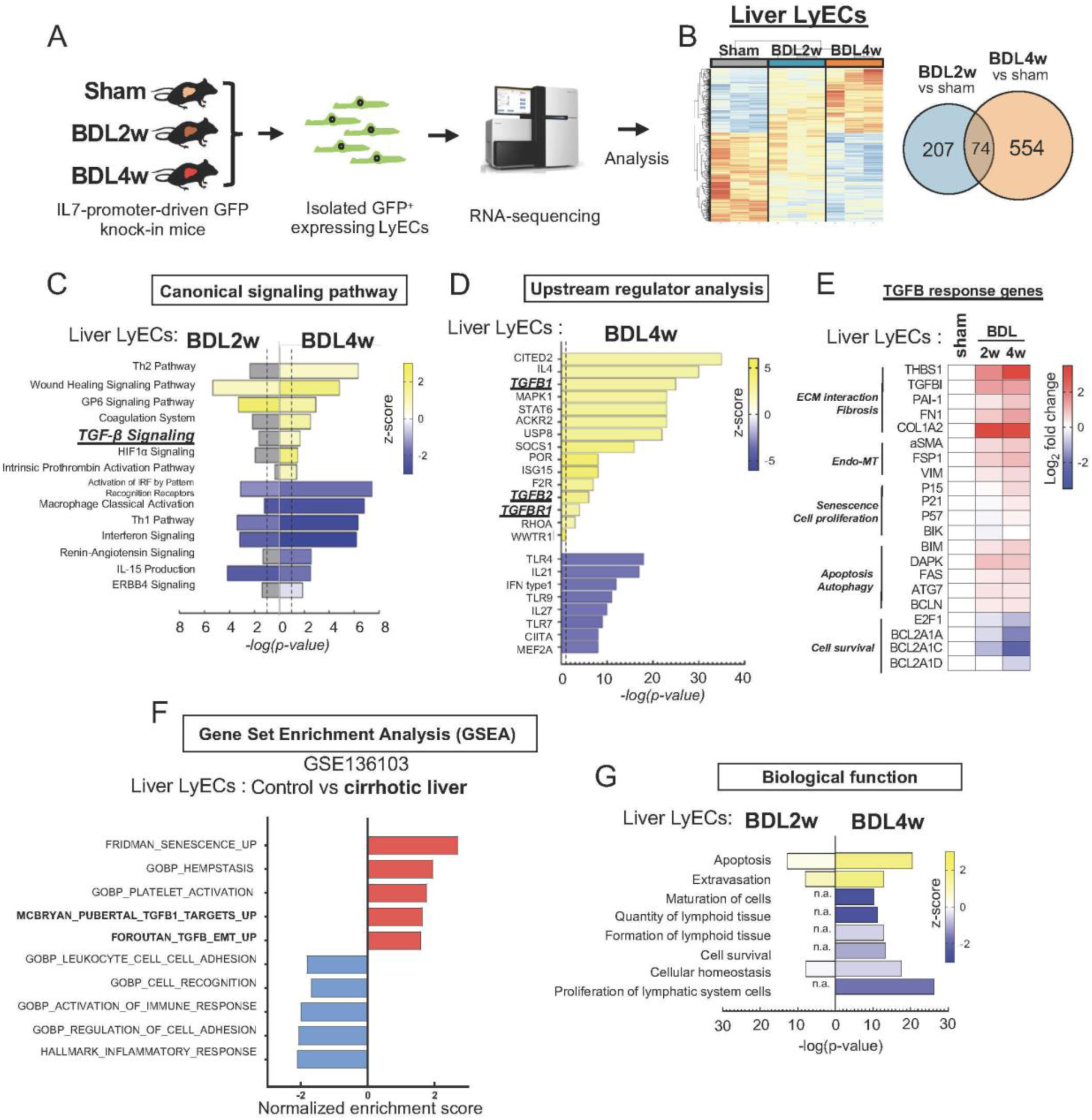
TGF-β signaling is up-regulated in lymphatic endothelial cells (LyECs) from cirrhotic human and mouse livers. (A) Schematic of liver LyECs isolation and its RNA sequencing in mice. RNA sequencing was performed in primary mouse liver LyECs isolated from IL7-promoter-driven GFP knock-in mice 2 weeks(n=3) and 4 weeks(n=3) after BDL or sham surgery (n=3). (B) Heatmap (left) and Venn diagram (right) showing genes differentially expressed in livers from BDL2w and BDL4w compared to Sham mice (fold change > 1.5; P ≤ 0.01). (C) Ingenuity Pathway Analysis (IPA) showing canonical signaling pathways altered in mouse liver LyECs from BDL2w(n=3) and BDL4w(n=3) mice. (D) Top potential upstream regulators in liver LyECs from BDL4w mice. (E) Heatmaps displaying TGF-β response genes differentially expressed in sham (n=3), BDL2w (n=3) and BDL4w (n=3). (F) The list of the Gene Set Enrichment Analysis(GSEA) signaling pathways altered in human liver LyECs (n=9) in cirrhotic patients from different etiologies compared to normal human subject (n=6). GSE136103. (G) The IPA biological functions altered in mouse liver LyECs from BDL2w(n=3) and BDL4w(n=3) mice. The vertical line indicates a Benjamini-Hochberg correction at −log, equating to P = 0.05.

Ingenuity Pathway Analysis (IPA) of the differentially expressed genes (DEGs) identified upregulation of TGF-β signaling in LyECs from BDL4w mice (**Figure 6C**). Specifically, TGF-β1/β2 and TGFBR1 were emerged as key upstream regulators in BDL4w LyECs (**Figure 6D**). Additionally, genes involved in TGF-β-mediated processes were upregulated, including those related to extracellular matrix (ECM) regulation (e.g., THBS1, TGFBI, and PAI-1), endothelial-to-mesenchymal transition (Endo-MT; e.g., aSMA, FSP1, and VIM), cellular senescence (e.g., P15, P21 and P57), and apoptosis (e.g., BIM, DAPK, FAS, and ATG7) (**Figure 6E**).

Analysis of a public dataset of human liver LyECs (GSE136103) corroborated these findings, showing upregulation of TGF-β signaling in LyECs from cirrhotic patients compared healthy controls (**Figure 6F**). Activation of TGF-β signaling is known to inhibit dermal LyEC proliferation and migration in inflamed environments(39, 40), which may contribute to the observed decrease in intrahepatic LVs in late-stage decompensated cirrhosis. Indeed, functional analyses revealed that “proliferation of lymphatic system cells”, “cell survival”, and “quantity of lymphoid tissue” were among the most downregulated biological processes in LyECs from BDL4w mice, suggesting TGF-β signaling as a central factor in LV regression during late-stage cirrhosis (**Figure 6G**).

### Blocking TGF-β1 signaling restores liver lymphatic vessel regression in late-stage decompensated liver cirrhosis

To test the hypothesis that enhanced TGF-β signaling in liver LyECs inhibits lymphangiogenesis and decrease LV numbers in decompensated cirrhotic livers, we first quantified TGF-β1 protein levels in livers from BDL mice. Livers from BDL4w mice exhibited the highest TGF-β1 levels (2.0-fold, p<0.05) compared to sham mice (**Figure 7A**). We next assessed Smad2 phosphorylation (pSmad2), a marker of TGF-β1 signaling activation. The number of pSmad2-positive LVs was significantly elevated in liver LyECs from BDL4w mice (∼5.6-fold increase, p<0.05) compared to sham, BDL1w and BDL2w mice (**Figure 7B**). Similarly, in human samples, a significant increase in pSmad2-positive LVs was observed in compensated cirrhotic patients, compared to healthy control, with further elevation in late-stage decompensated cirrhotic patients (**Figure 7C**). Additionally, the number of pSmad2-positive LV was inversely correlated with the total LV count in livers from compensated and decompensated cirrhotic patients (**Figure 7D**), indicating that TGFβ signaling negatively regulates LV numbers in cirrhotic livers.

**Figure 7.**
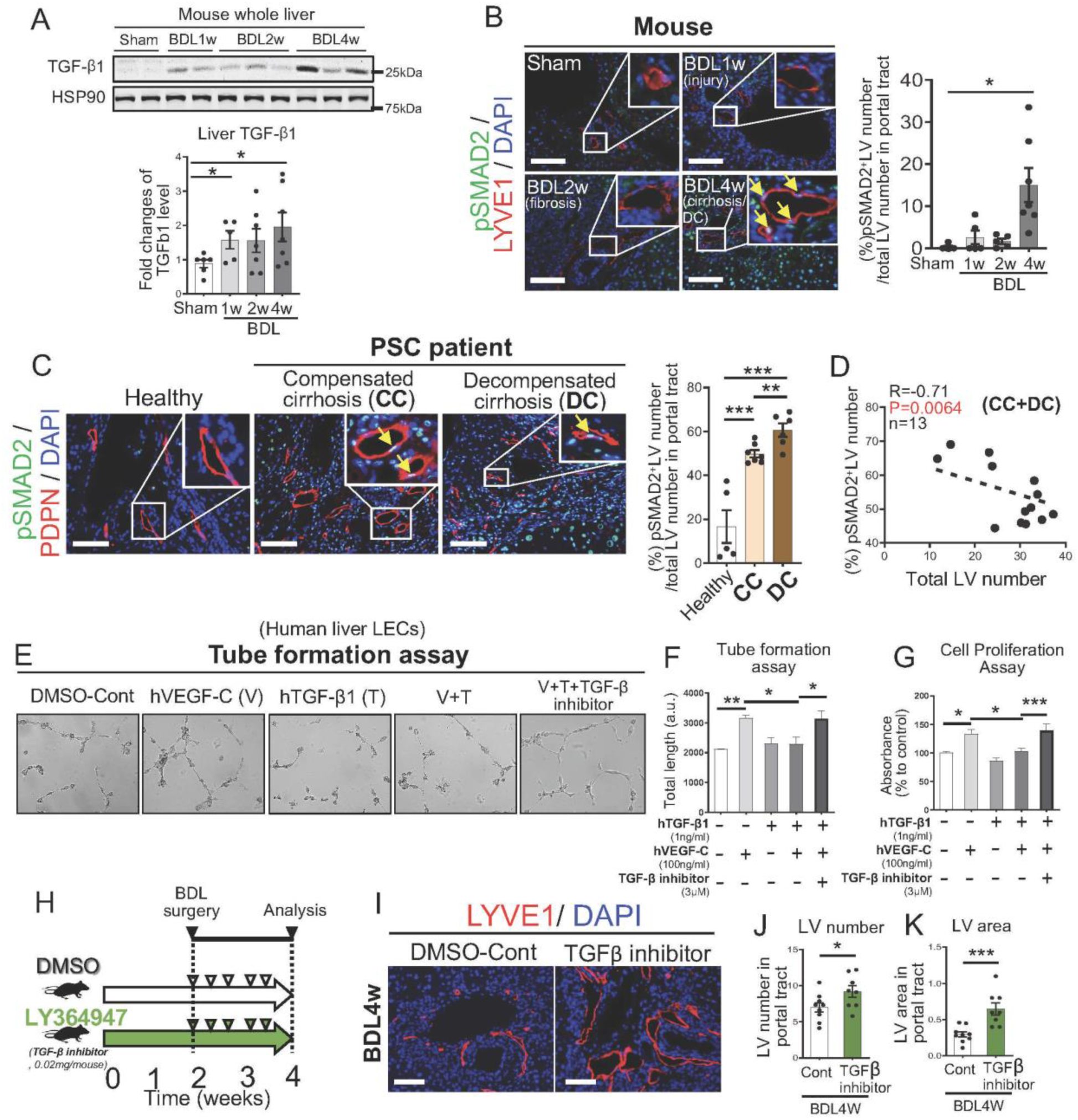
Increased TGF-β signaling is involved in regressed lymphatic vessel (LV) observed in decompensated cirrhosis in primary sclerosing cholangitis (PSC) patients and cholestatic mice. (A) TGF-β1 expression in livers from sham (n=6), 1-week(n=6), 2-week(n=5), and 4-week(n=7) BDL mice. (B) Representative immunofluorescence (IF) images of phospho-SMAD2 (green), LYVE1 (a LV marker; red) and DAPI in sham and 4-week BDL mouse livers. The arrows indicate Phospho-SMAD2-positive nuclei with DAPI in LYVE 1 positive lymphatic endothelial cells (LyECs). (right graph) Phospho-SMAD2-positive LVs number in livers from sham(n=5), 1-week(n=5), 2-week(n=5), and 4-week(n=7) BDL mice. (C) Representative IF images of phospho-SMAD2 (green), PDPN (red) with DAPI in healthy control livers(n=8) and PSC patients further classified into compensated cirrhosis (CC, n=7) and decompensated cirrhosis (DC, n=6). Quantification of phospho-SMAD2-positive LVs numbers in livers from healthy control (n=8) and PSC patients (CC vs. DC)(right panel). (D) A correlation between phospho-SMAD2-positive LV number and total LV number in PSC livers (n=13). (E) Representative images of tube formation assay to assess lymphangiogenesis and their quantification (F). Human primary liver LyECs incubated with or without VEGF-C (100ng/ml), hTGF-β1 (1ng/ml) and TGFβ inhibitor (LY364947) for 4 hours. Five images were taken per well and analyzed by the angiogenesis analyzer macro of ImageJ software. Representative data from three independent experiments are shown. (G) BrdU cell proliferation assay in human primary liver LyECs incubated with or without VEGF-C (100 ng/mL), hTGF-β1 (1ng/ml) and TGF-β inhibitor (LY364947) for 4 hours. Representative data from three independent experiments are shown. (H) Schematic of TGF-β inhibitor injection in mice. DMSO (control) or TGF-β inhibitor (LY364947, 0.02mg/mouse) was injected subcutaneously 2 weeks after BDL surgery followed by four more injections with a three-day interval between each administration until 4 weeks. (I) Representative immunofluorescence (IF) images and quantification (J, K) of LVs in livers from 4-week BDL mice treated with control-DMSO (n=8) or TGF-β inhibitor (n=8). LYVE-1 (a LV marker; red) and DAPI (blue). *P < 0.05, **P < 0.01, and ***P < 0.001. Scale bars: 100 μm.

We then explored the effects of TGF-β1 on the function of human primary liver LyECs. Treatment with human recombinant TGF-β1 (hTGF-β1) resulted in a time-dependent increase in pSmad2/3 signaling in liver LyECs (**Supplementary Figure 8A**). Furthermore, hTGF-β1 treatment upregulated TGF-β responsive genes, such as PAI1, while treatment with the selective ALK5 (TGF-β receptor 1) kinase inhibitor LY364947suppresed their transcription (**Supplementary Figure 8B**), confirming that liver LyECs respond to TGF-β1 signaling.

Functionally, TGF-β1 inhibited VEGF-C-induced tube formation (**Figure 7E, F**) and LyEC proliferation (**Figure 7G**). These inhibitory effects were reversed by TGF-β inhibitor LY364947 (**Figure 7E-G**). Together, these findings demonstrate that activation of TGF-β1 signaling reduces LV numbers and area in late-stage decompensated liver cirrhosis.

### Inhibition of endogenous TGF-β signaling restores liver lymphangiogenesis in late-stage cirrhosis in mice

To determine whether blocking TGF-β signaling can restore intrahepatic LV number and area in late-stage cirrhotic mice, we administered TGF-β inhibitor LY364947 starting 2 weeks after BDL surgery and continued treatment until 4 weeks post-surgery (**Figure 7H**). Livers from BDL4w mice treated with TGF-β inhibitor showed a significant increase in LV number and area compared to those untreated controls (**Figure 7I-K**). Furthermore, the median survival time of TGF-β inhibitor-treated BDL4w mice was extended by 3 days compared to control mice (**Supplementary Figure 9A**). However, TGF-β inhibition did not significantly reduce liver fibrosis as assessed by Sirius red staining and hydroxyproline content, nor did it alleviate liver injury, indicated by ALT levels (**Supplementary Figure 9 B-E**). Interestingly, blocking TGF-β signaling in BDL4w mice led to a reduction in fibrosis-related proteins, including collagen I (p=0.06), collagen III (p<0.05) and fibronectin (FN1) (p=0.07) (**Supplementary Figure 9 F&G**). These findings suggest that while inhibiting TGF-β signaling effectively restored LV numbers, this alone is insufficient to substantially resolve liver fibrosis in late-stage cirrhosis.

## DISCUSSION

This study demonstrates that impaired lymphatic drainage contributes to liver injury, fibrosis, and portal hypertension, driving the progression from compensated to decompensated cirrhosis. A key finding is the regression of lymphatic vessels (LVs) in late-stage cirrhosis, observed in both cholestatic mice and primary sclerosing cholangitis (PSC) patients. This LV regression, driven by elevated TGF-β signaling in lymphatic endothelial cells (LyECs), impairs lymphatic drainage, exacerbating inflammation and fibrosis. Conversely, liver-targeted VEGF-C overexpression enhances lymphatic drainage, reduces liver fibrosis and liver injury, and alleviates portal hypertension without affecting systemic hemodynamics. These results identified LV regression as a critical factor in liver decompensation and establish the therapeutic potential of restoring hepatic lymphatic function.

Using a surgical model to block liver lymphatic outflow, we provided direct evidence of the role of lymphatic dysfunction in liver pathology. Blocking lymphatic drainage increased hepatic stellate cell (HSC) activation, collagen deposition, and immune cell infiltration, underscoring the essential role of functional lymphatic drainage in maintaining liver health. One possible mechanism underlying lymphatic occlusion (LO)-induced fibrosis involves increased hydrostatic pressure due to excessive fluid (lymph) retention in the sinusoidal microcirculation and liver interstitial space, where HSCs reside. This is supported by increased spleen weight, a marker of increased intrahepatic resistance and hydrostatic pressure(41).

Elevated hydrostatic pressure has been shown to exacerbate fibrosis by activating mechanotransduction pathways that induce collagen alignment and myofibroblast differentiation(42–44), ultimately promoting fibrosis(42). In liver, increased hydrostatic pressure has been shown to induce the activation HSCs *in vitro*(42, 44). Our finding of increased liver and spleen weight, HSC activation (α-SMA expression), and sinusoidal fibrosis following LO surgery suggest that increased hydrostatic pressure contributes to the fibrogenic response in this model.

These results align with previous evidence that lymphangiogenesis enhances lymphatic drainage in liver disease models, particularly in fibrosis and cirrhosis(45, 46). While early-stage fibrosis is marked by compensatory lymphangiogenesis, our data reveal significant LV regression in late-stage cirrhosis in cholestatic mice and PSC patients. This regression correlates with impaired lymphatic drainage and the transition to decompensation. Supporting this, an intravital fluorescence microscopy study in rats with CCl_4_-induced liver fibrosis and cirrhosis reported increased functional LV density in early fibrosis (weeks 1–4), followed by a decline in late-stage fibrosis (weeks 8–12)(1). The regression of intrahepatic LV in late decompensated cirrhosis we observed in our study could partially explain the defect in lymphatic drainage function seen in patients(46, 47) and pigs(47) with late decompensated cirrhosis stage. Collectively, these findings suggest that reduced LV numbers and impaired drainage contribute to the progression from compensated to decompensated cirrhosis.

We identified TGF-β as a central driver of LV regression in late-stage cirrhosis. Previous studies have shown that TGF-β suppresses LyEC proliferation, migration, and tube formation(39, 40, 48). While TGF-β receptor I (TGFβRI) inhibitors restore LV numbers in mouse models of traumatic injury, chronic peritonitis, and pancreatic cancer(39, 49), they are insufficient to fully prevent fibrosis or lymphedema in an injured tail assay(50). Interestingly, the effect of TGF-β1 inhibition on lymphangiogenesis is enhanced in the presence of VEGF-C(39), suggesting a synergistic approach combining VEGF-C with TGF-β1 inhibitors may effectively restore lymphatic function in late-stage cirrhosis. VEGF-C has been shown to mitigate TGF-β’s anti-lymphangiogenic effects and restore lymphatic function at higher concentrations(51).

In this study, liver-targeted VEGF-C overexpression significantly restored lymphatic drainage, reduced fibrosis, alleviated portal hypertension, and mitigated liver injury. We propose that for treating late-stage cirrhosis, AAV8-mediated prolonged VEGF-C expression may be a promising approach. Unlike recombinant VEGF-C (Cys156Ser) formulations used in a rat study(29), which primarily targets mesenteric lymphatics, AAV8-VEGF-C delivery focuses on intrahepatic lymphatics, leveraging the liver-specific tropism of the AAV8 vector(52). A study in a pig model of lymphedema demonstrated that VEGF-C provides superior lymphangiogenic effects with minimal vascular side effects compared to the Cys156Ser form of VEGF-C(53). Our liver-targeted strategy addresses the root causes of portal hypertension and fibrosis, while avoiding systemic hemodynamic effects, such as splanchnic vasodilation.

Importantly, VEGF-C activates endothelial nitric oxide synthase (eNOS)(54), improving hepatic microcirculation and reducing vascular resistance. Since LSECs express VEGFR3(55), VEGF-C overexpression may ameliorate LSEC dysfunction and capillarization, enhancing liver lymphatic drainage, as plasma filtration through sinusoidal microcirculation is the first step in lymph production. Our observation of decreased CD34 expression in LSECs in cirrhotic mouse liver as a result of VEGF-C overexpression (Suppl Fig 5) support this possibility.

In conclusion, this study highlights the crucial role of liver lymphatic drainage in maintaining liver homeostasis and regulating disease progression. We identify LV regression and impaired lymphatic function, mediated at least in part by TGF-β elevation, as central mechanisms driving liver decompensation. Furthermore, our findings position VEGF-C overexpression as a promising therapeutic strategy to restore hepatic lymphatic function, reduce fibrosis and alleviate portal hypertension. These insights provide a foundation for the development of lymphatic-targeted therapies with broad applicability across liver diseases.

## MATERIALS AND METHODS

### Study approval

All animal experiments were approved by the Institutional Animal Care and Use Committee of the Veterans Affairs Connecticut Healthcare System and by the Taipei Veterans General Hospital Animal Committee (IACUC 2023-130), and conducted in accordance to the guidelines for laboratory animal care as specified in the National Institutes of Health Guide for the Care and Use of Laboratory Animals (DHEW publication No. (NIH) 85-23, rev. 985, Office of Science and Health Reports, DRR/NIH, Bethesda, MD, USA).

### Sex as a biological variable

Our study exclusively examined male mice. It is unknown whether the findings are relevant for female mice.

### Liver lymphatic occlusion (LO) surgery

Male 6- to 10-weeks-old, C57BL/6 mice were used for liver lymphatic occlusion (LO) surgery. LO or sham surgery was performed under anesthesia with ketamine (100 mg/kg) and xylazine (10 mg/kg). Briefly, the mice’s abdomen and flank areas were shaved and disinfected with povidone-iodine. A 2-cm incision was made in the abdominal area, and the portal lymph node (pLN) and celiac lymph node (cLN), where the afferent lymphatic vessels from the liver run into, were identified under the stereomicroscope. After removing these tow two lymph nodes (cLN& pLN), in order to complete block the lymphatic drainage, 5-10 μl Gorilla Super Glue Micro Precise Gel was applied to the area where the celiac LN and portal LN were located. Liver samples were collected at 3 days after LO surgery for analysis. In sham mice, the same surgery except for the LO procedure was performed.

### Bile duct ligation (BDL) surgery for inducing cholestasis in mice

Male 6- to 10-weeks-old, tamoxifen administrated endothelial-specific green fluorescent protein-reporter (EC-GFP reporter) mice underwent BDL surgery unless otherwise mentioned. BDL or sham (control) surgery was performed under anesthesia with ketamine (100 mg/kg) and xylazine (10 mg/kg)(24). Liver samples were collected at 1, 2 and 4 weeks after BDL surgery. In sham mice, the same surgery except for the ligation procedure was performed, and liver samples were collected 1 week after BDL surgery.

EC-GFP reporter mice were generated by crossing Cdh5-CreERT2 mice(56) and ROSA^mTmG^ mice (Strain#:007676, The Jackson Laboratory, Bar Harbor, ME). For cre-recombination, EC-GFP reporter mice were injected with tamoxifen (100 mg/kg) (Cat# T5648, Sigma-Aldrich) dissolved in corn oil intraperitoneally for 5 consecutive days at least 3 weeks before experiment.

### Manipulation of the VEGF-C/VEGFR3 signaling in cholestatic mice

To examine the effect of enhanced lymphangiogenesis by VEGF-C overexpression, mice were randomly divided into two groups, one given Adeno-Associated Virus (AAV)8-mouse VEGF-C (Vector Biolabs, Malvern, PA) and the other given AAV-GFP (control) (Cat#37825-AAV8, Addgene, Watertown, MA). AAV-VEGF-C and AAV-GFP were prepared in Dulbecco’s Phosphate-Buffered Saline (Cat#14190-144, DPBS, Gibco-Thermo Fisher Scientific, Waltham, MA) with a concentration of 2.0 x 10^10^ GC/ mouse in a total volume of 100μl on the day of injection, and injected in the retroorbital region one week before BDL surgery. Livers and biological samples were collected 2 weeks after BDL surgery (i.e., 3 weeks after AAVs injection) for further analysis.

To examine effect of blocking the lymphangiogenesis, MAZ51 (a selective VEGFR-3 Tyrosine Kinase Inhibitor, Cat#676492, Millipore Sigma, Billerica, MA), was used, Immediately after BDL surgery, mice were randomly divided into two groups, one given MAZ51 and the other given DPBS containing 20% of Dimethyl Sulfoxide (DMSO; Cat#276855, Sigma-Aldrich) (control). MAZ51 solution was freshly prepared in DPBS containing 20% of DMSO with a concentration of 2mg/ml. MAZ51 solution in 100μl (0.2mg/mouse) was injected Intraperitoneally right after BDL surgery. Similarly, equal volume (100μl) of control DPBS solution containing 20% DMSO was injected just after BDL surgery as control. Subsequently, we repeated the MAZ51 and control DPBS solution injections two more times, with a three-day interval. Livers and biological samples were collected one week after BDL surgery for further analysis.

### Bile duct ligation (BDL) surgery for inducing cholestasis in rats

Sprague-Dawley rats, initially weighing between 240 and 270g, were used for the experiments. During the study, the rats had unlimited access to food and water. Secondary biliary cirrhosis was induced by ligating the common bile duct. Anesthesia was provided by administering Zoletil (a 1:1 mixture of tiletamine and zolazepam, Virbac, Carros, France) intramuscularly at a dose of 50 mg/kg. The surgical procedure involved placing two ligatures using 3-0 silk: one below the hepatic duct junction and the other above the pancreatic duct entry. The common bile duct was then cut between these ligatures. Following either bile duct ligation or a sham operation, AAV8-GFP (1.5 × 10^12^ cp/rat, AAV Core Facility of Academia Sinica, Taipei, Taiwan) or AAV8-VEGFC (1.5 × 10^12^ cp/rat, AAV Core Facility of Academia Sinica, Taipei, Taiwan) was administered via the tail vein. Postoperative care was then provided. To prevent coagulation issues, rats received weekly intramuscular injections of vitamin K at a dose of 50 μg/kg following the ligation. The hemodynamic measurements were performed 4 weeks after operation.

### Systemic and portal hemodynamics measurements

Hemodynamic measurements were conducted according to protocols established in a previous study(57). Briefly, a PE-50 catheter was inserted into the right carotid artery and connected to a Spectramed DTX transducer (Spectramed Inc., Oxnard, CA, USA). Using a multi-channel recorder (MP45, Biopac Systems Inc., Goleta, CA, USA), continuous recordings of mean arterial pressure (MAP), heart rate (HR), and portal pressure (PP) were obtained. An external zero reference was set at the midpoint of the rat’s body. An abdominal midline incision was then made, and the mesenteric vein was cannulated with another PE-50 catheter connected to a transducer.

The superior mesenteric artery (SMA) was identified at its origin from the aorta, and a 5-mm segment was carefully isolated. A pulsed-Doppler flow transducer (TS420, Transonic Systems Inc., Ithaca, NY, USA) was positioned to monitor SMA flow. Portal flow was measured using an appropriately sized flow transducer placed near the liver.

Cardiac output (CO) was measured using the thermodilution technique, involving a thermistor placed in the aortic arch just beyond the aortic valve and injecting 100μl of normal saline into the right atrium via a PE-50 catheter. The Cardiomax III cardiac output computer (Columbus Instruments International Co., OH, USA) processed the thermodilution curves, with five curves obtained for each measurement. The final CO value was the average of these results. Cardiac index (CI, ml/min/100g BW) was calculated by dividing CO by the body weight (BW) in grams. Systemic vascular resistance (SVR, mmHg/ml/min/100g BW) was determined by dividing MAP by CI. Superior mesenteric artery resistance (mmHg/ml/min/100g BW) was calculated using the formula (MAP - PP)/SMA flow per 100g BW. Hepatic vascular resistance (HVR, mmHg/ml/min/100g BW) was derived by dividing PP by hepatic inflow (portal portion) per 100g BW.

### Human liver specimens

Thirteen liver specimens from patients with Primary Sclerosing Cholangitis (PSC; n=13) and healthy controls (n= 8) without liver disease were used. These liver specimens were acquired from the Liver Tissue Cell Distribution System at the University of Minnesota (funded by National Institutes of Health Contract # HSN276201200017C). These specimens are also coded without patient/donor identities. We did not ascertain individual identities associated with the specimens. The patients ranged in age from 25 to 68 years old (mean 47.1 y) and included 13 males. We categorized hepatic compensated cirrhosis (CC) and decompensated cirrhosis (DC) based on a previous study(38). History of hepatic DC (n= 6) was defined as evidence of prior diagnosis of ascites, hepatic encephalopathy, variceal hemorrhage, or baseline concomitant medications with a specific indication for ascites, hepatic encephalopathy, or variceal hemorrhage, while CC patients (n= 7) were those with absence of complications listed above for hepatic DC patients. The clinical information of these patients were summarized in Supplementary Table 2. All liver specimens were fixed in 10% neutral buffered formalin and embedded in paraffin (paraffin blocks) for histological analysis.

### Immunohistochemistry (IHC) for PDPN-positive human liver lymphatic vasculature

Human liver paraffin sections at a thickness of 7μm were deparaffinized with xylenes and rehydrated by washing through a graded alcohol series to deionized water. followed by washing in PBS 3 times. For antigens retrieval, the sections were incubated with 10mM citrate buffer (pH 6.0) for 15 minutes at approximately 100°C. The sections were then cooled down for 20 minutes, washed in PBS 3 times, incubated with 3% H_2_O_2_ solution dissolved in MeOH for 30 min at room temperature to block peroxidase activity, then washed with PBS 3 times. To block non-specific binding of antibodies, the sections were incubated with normal horse serum blocking buffer (Cat#PK-6102, VECTASTAIN® Elite® ABC-HRP Kit, Vector Laboratories, Burlingame, CA) for 1 hour at room temperature and briefly washed with PBS. Additionally, the sections were incubated with Avidin D solution for 15 minutes followed by Biotin solution for 15 minutes at room temperature using Avidin/ Biotin Blocking Kit (Cat#SP-2001, Vector Laboratories), and briefly washed with PBS. For PDPN staining, the sections were incubated with 70 μl of primary PDPN antibody (ready to use, Cat# M3619, Dako, Carpinteria, CA) at 4 °C, overnight, and washed in PBS 3 times. The sections were then incubated with biotinylated anti-mouse IgG secondary antibody (VECTASTAIN® Elite® ABC-HRP Kit, Vector Laboratories) for 30 minutes, washed in PBS 3 times followed by applying avidin-biotin-peroxidase complex (VECTASTAIN® Elite® ABC-HRP Kit, Vector Laboratories) for 30 minutes at room temperature, and washed again with PBS 3 times. The PDPN staining was visualized as brown color by reaction of the coupled peroxidase with 3,3′-diaminobenzidine (Cat#SK-4100, DAB Peroxidase Substrate Kit, Vector Laboratories), as recommended by the manufacturer. The stained sections were washed in PBS 3 times, and stained with Meyer’s hematoxylin counterstaining solution (Cat#MHS32, Sigma-Aldrich, St. Louis, MO) for 15 sec followed by washing in warm tap water for 5 min. Paraffin sections were dehydrated by washing through deionized water to graded alcohol series and Xylene. The dehydrated stained sections were mounted with histology mounting medium (Cat#8310-4, Cytoseal ™ 60, Thermo Fisher Scientific, Waltham, MA).

### Inhibition of TGF**-**β signaling in BDL mice

Mice at 2-week BDL surgery were randomly divided into two groups, one given TGF-β inhibitor, a TGF-β Type I Receptor Kinase Inhibitor (LY364947, Cat#L6293, Sigma-Aldrich) and the other given DMSO (control). LY364947 solution was prepared in DPBS containing 8% of DMSO with a concentration of 0.2mg/ml on the day of injection. 100 μl (0.02mg/mouse) of LY364947 solutions were then injected intraperitoneally 2 weeks after BDL surgery. As a control, equal volume (100 μl) of DPBS solution containing 8% DMSO was injected 2 weeks after BDL surgery. Subsequently, we repeated the injections, either LY364947 or control solution five more times, with a three-day interval between each administration. Livers and biological samples were collected 4 weeks after BDL surgery (i.e., 2 weeks after LY364947 injection) for further analysis.

### Immunofluorescence (IF) staining

#### IF for frozen section of mouse livers

Mouse livers were fixed in 4% paraformaldehyde for 48 hours at 4 °C, which were frozen in a mold containing O.C.T compound (Cat#4583, Sakura, Miles, Elkhart, IN). Frozen blocks were cut at a thickness of 7 μm and these liver sections were washed in PBS for 10 minutes 3 times. To retrieve antigens, the sections were incubated with 10 mM citrate buffer (pH 6.0) for 15 minutes at approximately 100 °C. In the case of CD3, CD19, MPO immunolabeling, the antigens retrieval procedure was not performed. Detail staining conditions are provided in Supplementary Table1. After cooling down for 20 minutes, sections were washed in PBS 2 times (most internal reporter fluorescent signals were removed after retrieving antigens procedure). Then, the sections were incubated with 5% donkey serum solution containing 0.3% Triton X-100 for 1 hour at room temperature (blocking process), briefly washed in PBS, and incubated with primary antibodies [LYVE1 (dilution factor: 1:300, Cat#AF2125, R&D systems, Minneapolis, MN), α-SMA (1:300, Cat# NB300-978, NOVUS Biologicals, Centennial, CO), CD68 (1:100, Cat# MCA1957, Blackthorn, Oxon, UK), CD3 (1:100, Cat# MAB4841, R&D systems), MPO (read to use, Cat# PP023AA, Biocare Medical, Concord, CA), CD19 (1:100, Cat# NBP2-15782, NOVUS Biologicals), Albumin (ALB, 1:50, Cat# sc-46289, Santa Cruz Biotechnology, Santa Cruz, CA), CD34 (1:100, Cat#553731, BD Biosciences, Oxford, UK), Desmin (1:100, Cat#5332, Cell Signaling Technology Danvers, MA), pSMAD2 (1:100, Cat#3102, Cell Signaling Technology), PCNA (1:100, Cat#2586, Cell Signaling Technology)], dissolved in 5% donkey serum solution containing 0.3% Triton, at 4 °C, overnight. After washing with PBS 3 times, the sections were incubated with secondary antibody solution, including donkey anti-rabbit Alexa Fluor 647 conjugate, anti-mouse Alexa Fluor 647 conjugate, anti-goat Alexa Fluor 488 conjugate, donkey anti-rat Alexa Fluor 647 conjugate, or donkey anti-rat Alexa Fluor 488 conjugate with 1:300 dilution for 30 minutes at room temperature. After washing in PBS 3 times, mounted with Fluoroshield™ with DAPI histology mounting medium (Cat#F6057, Sigma-Aldrich).

#### IF for paraffin sections for mouse and human livers

mouse and human livers were fixed in 10% neutralized buffered formaldehyde for 48 hours at 4 °C. The paraffin blocks were prepared and cut at a thickness of 7 μm, and subsequent deparaffinization, antigen retrieval and blocking procedures were as described above in the IHC section. The liver sections were incubated with primary antibodies for mouse livers [LYVE1 (1:200, Cat#AF2125, abcam, Minneapolis, MN), CK19 (1:50, Cat#TROMA-III, DSHB, Iowa City, IA)], or human livers [PDPN (ready to use, Cat# M3619, Dako), pSMAD2 (1:100, Cat#3102, Cell Signaling Technology)] at 4 °C, overnight. The sections were then washed with PBS 3 times, and were incubated with secondary donkey anti-rabbit Alexa Fluor 647 conjugate, donkey anti-rabbit Alexa Fluor 647 conjugate, donkey anti-rat Alexa Fluor 488, or anti-mouse Alexa Fluor 647 conjugate for 30 minutes at room temperature. All secondary antibodies dilution factor was 1:300. The sections were washed with PBS 3 times and mounted as described above.

### Sirius Red staining and quantification

Sirius Red staining was performed on formalin-fixed paraffin-embedded liver tissues. Paraffin sections (7 µm thickness) were deparaffinized and rehydrated as described above, and then stained with picro-sirius red (F3B, Direct Red 80, Cat#365548, Sigma-Aldrich) for 1 hour, and washed by 0.5% acetic acid (glacial) solution 2 times, followed by dehydration with increasing concentrations of ethanol and clearing with xylene. The dehydrated stained sections were mounted with histology mounting medium (Cat#8310-4, Cytoseal ™ 60, Thermo Fisher Scientific). For the quantification of Sirius Red positive-fibrotic area, the images were obtained using the Olympus BX51(Olympus, Tokyo, Japan) microscope system with a 20 x objective lens. Five to ten images of the liver portal tract area were taken per liver section slide. Sirius Red-positive area was quantified using the ‘polygon selection’ tool in Image J [National Institutes of Health (NIH), Bethesda, MD].

Quantification of PDPN-positive lymphatic vessel (LV) in human liver. First the IHC images of PDPN-positive LVs were taken using the Olympus BX51 (Olympus) microscope system with a 10 x objective lens. Five to ten images of the liver portal tract area were taken per liver section slide. Image J (NIH) was used for quantification of the PDPN-positive LV (luminal) area in human livers. Briefly, the liver LV number and area were captured and measured using the ‘polygon selection’ tool in the Image J manually. We have excluded PDPN-positive bile duct based on cuboidal epithelial structure. For LYVE1-positive LVs or αSMA-positive vessels, such as hepatic artery (HA) and portal vein (PV) quantifications in mouse liver, the IF images were taken using the Zeiss Axio observer microscope system (Carl Zeiss, Munich, Germany) with a 20x objective lens. The Zeiss Axio observer microscope system with a 20x objective lens was used for all the image acquisition in IF stained liver section slides unless otherwise mentioned. Five to ten images of the liver portal tract area were taken per liver section slide. Image J was used for quantification of the LYVE1-positive LV number and area and αSMA-positive HA number/area, or PV area in mouse livers as described above. *Percentage (%) of PCNA-positive proliferating LV number was determined by taking*, five to ten images of the portal area per liver section slide, and divided by total LYVE1-postive LVs number. PCNA and DAPI double-positive nuclei in the lining of LYVE 1-positive LVs were counted as proliferating LV. Similarly, for assessing % of PCNA-positive proliferating hepatocyte quantification in mouse livers, five to ten images of the mid-lobular zone 2 area(58) were taken per liver section slide, and PCNA and DAPI double-positive nuclei in albumin (ALB)-positive hepatocytes were counted as proliferating hepatocyte. Then, % of PCNA-positive proliferating hepatocytes were calculated by dividing total DAPI-positive nuclei number in ALB-positive area. For % of pSMAD2-positive LVs number quantification in mouse/human livers by IF staining, five to ten images of the portal tract area were taken per liver section slide. pSMAD2 and DAPI double-positive nuclei in the lining of LYVE1 or PDPN-positive LVs were counted as pSMAD2-positive LV. Percentage of pSMAD2-positive LVs number were calculated divided by total LYVE1 or PDPN-positive LVs number in PT area. For the quantification of CD68-, CD3-, MPO-, CD19-positive immune cells in mouse liver by IF staining, five to ten images of the portal tract area were taken per liver section slide. Image J (NIH) was used for quantification of the CD68-, CD3-, MPO-, CD19-positive macrophage, T cell, neutrophil, and B cell number in mouse livers. The immune cells number were captured and counted in designated portal tract area using the ‘polygon selection’ tool in the Image J. For the quantification of CK19-positive bile duct (BD) in mouse liver, five to ten images of the portal tract area were taken per liver section slide. The BD area was measured in the designated portal tract area by the ‘polygon selection’ tool in Image J. For the quantification of desmin/αSMA double-positive activated hepatic stellate cell (HSC), five to ten images of the portal tract area were taken per liver section slide. For quantification of CD34-positive LSECs, five to ten images of the liver lobular zone2-3 area(35) were taken per liver section slide, and the CD34-positive area was measured in the entire image using Image J.

### Liver hydroxyproline level measurement

Frozen liver samples (−80 °C) (between 45 and 55 mg) was homogenized in a 1.5 ml tube containing of 6N HCl solution (10 µl/mg tissue) using a pestle. The homogenate was incubated for 24h at 110°C. After brief centrifugation and cooling down at room temperature, the homogenate was filtered using a 0.45-micron syringe filter. 25 µl of filtered homogenate was transferred to a new 1.5ml tube and added to 225 µl of 2.2% NaOH solution [2.2 g Sodium hydroxide dissolved in 100ml of citrate-acetate-buffer (5 g Citric acid monohydrate · 1.2 ml Acetic acid · 12 g Sodium acetate trihydrate · 3.4 g Sodium hydroxide dissolved in H_2_O to a final volume of 100ml, and adjusted pH 6.0)]. 125 µl of Chloramine-T-solution [0.141g Chloramine-T hydrate (Cat# 857319, Sigma-Aldrich) · 2ml H_2_O · 3ml 2-Methoxyethanol (Cat#E5378, Sigma-Aldrich) · 5ml citrate-acetate-buffer] was added to neutralized homogenate and incubated for 20 min at room temperature. Subsequently, 125 µl of Perchloric acid (Cat#100519, Millipore Sigma) was added and incubated for 20min at room temperature, followed by adding 125 µl of dimethyl benzaldehyde solution [2g 4-(Dimethylamino)benzaldehyde (Cat#D2004, Sigma-Aldrich) dissolved in 10 ml of 2-Methoxyethanol (Sigma-Aldrich)] and incubation for 20 min at 60°C. After incubation, the absorbance of the final mixture was determined at 565nm. For standard solutions, trans-4-Hydroxy-L-proline (Cat#H54409, Sigma-Aldrich) was dissolved in citrate-acetate-buffer concentration with 6 µg /ml (working solution). To generate the standard curve, each standard solution containing 6, 3, 1.5, 0.75, 0.375, 0.1875, 0.09375 and 0 µg/mL of trans-4-Hydroxy-L-proline was prepared with serial dilution of working solution in citrate-acetate-buffer. 250 μl of each standard solution was added 125 µl of Chloramine-T-solution and treated according to the above procedure for the following step. The value of the liver hydroxyproline level was expressed as µg/g wet tissue.

### Alanine Transaminase (ALT) measurement

Mouse plasma ALT levels were determined using the ALT kit (Cat# A7526-150, Pointe Scientific, Canton, MI) according to the manufacturer’s instructions. Blood samples were collected from the vena cava into heparin-coated collection tubes, and plasma was separated from blood after centrifugation with 2800 rpm for 20 min at 4°C.

### Liver lymphatic drainage function test

The liver lymphatic drainage function was performed with some modifications of previous publications(59, 60). Evans blue working solution was prepared as one percent Evans Blue dye (Cat#E2129, Sigma–Aldrich) dissolved in Hank’s Buffered Salt Solution (Cat# 14175095, Gibco-Thermo Fisher Scientific, Waltham, MA). 10 µl of Evans blue working solution was injected directly into exposed liver parenchyma (right, left, and median lobes) using a 27-gauge 1/2 syringe in mice under anesthesia with ketamine (100 mg/kg) and xylazine (10 mg/kg) in mice, and kept for 10 min in mice under continuous anesthesia to allow the Evans blue dye to migrate through the LVs and reach liver draining portal lymph node (portal LN), and then euthanized. Portal LNs were immediately collected using a dissecting scissors/forceps. Evans blue dye was extracted from portal LNs in a 1.5 ml tube containing of 300 μl of N, N-dimethyl formamide (Cat#68-12-2, Millipore Sigma), which was homogenized using a pestle and incubated overnight at 55°C. The following day, the Evans blue content from individual Portal LN samples was measured with an excitation at 620 nm and an emission of 680 nm, and determined using a standard curve generated by 10, 5, 2.5, 1.25, 0.625, 0.3125, and 0 µg/mL of Evan blue, prepared by serial dilution in N, N-dimethylformamide (Millipore Sigma).

### Primary mouse liver lymphatic endothelial (LyEC) cell isolation

Recently, we have identified that IL7 is exclusively expressed by LyEC in the liver, not by other liver cells(61). Primary mouse liver LyECs were isolated from IL7GFP/+ knock-in mice at 2 weeks and 4 weeks after BDL surgery or sham mice using our established protocol(61). Briefly, the liver was digested by warm collagenase IV (Cat# 07427, STEMCELL Technologies, Vancouver, CA) perfusion (0.5 mg/ml, 37°C) through the portal vein under anesthesia. The perfused liver was isolated and transferred into a cell strainer in a 10cm petri dish containing RPMI 1640 (Cat#11875-093, Gibco-Thermo, Waltham, MA) with 10% fetal bovine serum (FBS) (Cat#16140-071, Gibco-Thermo). Digested liver cells were dispersed by gently rubbing with a syringe plunger, only leaving an undigested portal vein (PV) tree portion containing the LVs, PVs, hepatic artery, and bile ducts in the cell strainer.

Liver LyECs were isolated from the PV tree, which was first transferred into a 10cm petri dish containing cold DPBS, carefully removed residual parenchymal cells using surgical forceps, and washed thoroughly again in a 60mm petri dish containing cold PBS. After washing, the PV tree was cut into small pieces with scissors, transferred into a 15ml conical tube, and centrifuged at 50 g for 3 minutes at room temperature. The supernatant was discarded, and the cell pellets were resuspended in 3 ml of enzymatic dissociation buffer [1x HBSS (Cat# 14175095, Gibco-Thermo) containing 5 mg collagenase IV and 360U DNase I (Cat#07470, STEMCELL Technologies)] and incubated for 1 hour 30 minutes at 37°C to dissociate cells. After incubation, 2.5 ml of DMEM (Cat#11965-092, Gibco-Thermo) containing 3% FBS was added, and cell suspension was transferred into a 10ml syringe equipped with a 21G needle to gently triturate in a 10cm petri dish. After repeating the trituration process using a 22G and then a 27G needle, the triturated cell suspension was collected and centrifuged at 400 g for 3 minutes at room temperature. The pellet was resuspended in 1x HBSS, triturated gently once again with a 27G needle, and centrifuged at 400 g for 3 minutes at room temperature to get a PV tree cell fraction. The PV tree cell fraction was used for LyEC isolation based on their GFP expression by FACS [the BD FACSAria IIu Cell Sorter (BD Biosciences, San Jose, CA)] using a 100μm nozzle. Isolated LyECs were used for RNA-seq analysis.

### RNA sequencing analysis of isolated liver LyECs

2×10^4^ to 1.6×10^5^ mouse liver primary LyECs were sorted and directly dissolved into 500 µl of TRIzol (Cat#15596-018, Invitrogen, Carlsbad, CA) reagent and total RNA was extracted as previously described(61). RNA samples were submitted to the Yale Center for Genomic Analysis, and libraries were constructed according to their protocols. The libraries underwent 100-bp paired-end sequencing using a NovaSeq instrument. Reads were trimmed using Trim Galore (v0.5.0) and mapped to the mouse reference genome (mm10) using HISAT2 (v2.1.0)(62). Gene expression levels were quantified using StringTie (v1.3.3b)(63) with gene models (v27) from the GENCODE project. Differentially expressed genes (DEGs) were identified using DESeq2 (v1.22.1)(64). Genes with a fold change of more than 1.5 or less than 2/3 as well as an adjusted p-value of less than 0.1 were considered as differentially expressed. A heatmap was generated using DEGs to illustrate gene expression changes in association with BDL injury. DEGs were also used for further functional comparisons between sham vs. BDL2w, sham vs. BDL4w and BDL2w vs. BDL4w groups using Ingenuity Pathway Analysis (IPA, Qiagen, Hilden, Germany) to create figures of the enriched canonical pathways, upstream regulator predictions, and biological function for each comparison.

### Human/Mouse liver tissue transcriptomic data analysis

All analyses were performed in the Seurat R package (version 4.3.0). For mouse liver scRNA-seq analysis, we retrieved our previously published data (GSE147581)(35) on endothelial cells (ECs) isolated from livers of control or fibrotic EC-specific GFP reporter mice (Cdh5-Cre-mTmG+/+ mice). Each cell cluster including LyEC, liver sinusoidal endothelial cell (LSEC), Venous endothelial cell (VEC), Arterial endothelial cell (AEC), Hepatic stellate cell (HSC), Cholangiocyte, B cell, T cell, Macrophage (MP), and hepatocyte from mouse liver scRNA-seq were identified and subsetted according to the criteria in previous study(61).

For human liver scRNA-seq analysis, a previously published human scRNA-seq data was downloaded from Gene Expression Omnibus (GEO) under an accession number GSE13610342(65). Cell clusters were annotated according to the original study. LyEC cluster was identified by expressing marker genes including CCL21, PDPN, and TFF3. Differential gene expressions (DEGs) analysis was conducted on LyEC cluster cells between healthy and cirrhotic patient samples using FindMarkers function in Seurat. Genes with a fold change of more than 1.5 or less than 2/3 as well as an adjusted p-value of less than 0.1 were considered as differentially expressed. Gene Set Enrichment Analysis (GSEA) (version 4.2.3)(66) was performed on DESeq2-normalized gene expression counts. Mouse ENSEMBL gene identifiers were mapped to human orthologs using the corresponding MSigDB.v7.5.1.chip. We set 1000-time permutations for all pathways.

### Cell culture conditions for human primay liver lyECs and

TGF-β treatment Human liver LyECs were purchased from Angio-Proteomie (Cat# cAP-0038, Lot# 201512302, Angio-Proteomie, Boston, MA), and sub-cultured with ENDO-Growth medium (Cat# cAP-05, Angio-Proteomie) on cell culture dish coated with Quick Coating Solution (Cat# cAP-01, Angio-Proteomie), cultured at 37°C, 5% CO_2_ in a humidified incubator. The human liver LyEC was incubated with basal medium (Cat#cAP-03, Angio-Proteomie) in the presence recombinant human TGF-β (hTGF-β, 1ng/ml) (Cat# 240-B, Bio-Techne/ R&D Systems) for 5min, 10min, 30min, 1hr, and 2hr. The control group was treated with an equal amount of DMSO, the same as the hTGF-β group for 2 hours. Total protein was collected for Western blot analysis. In addition, human liver LyEC was incubated with ENDO-Growth medium with or without TGF-β inhibitor (3uM) (LY364947, Cat#L6293, Sigma-Aldrich) in the presence with hTGF-β 1(1ng/ml) for 24h, then total RNA was collected for quantitative real-time polymerase chain reaction.

### Lymphatic endothelial cell tube formation assay

The ability of human liver LyECs to form capillary-like structures was examined with slight modifications from previously described(24). In brief, 10 mg/ml Matrigel® growth factor-reduced basement membrane matrix (Cat# 354230, Corning, Corning, NY) was loaded to a 24-well plate with 280 μl each. The plate was incubated at 37°C for 30 minutes. Human liver LyECs suspension with basal medium (Cat#cAP-03, Angio-Proteomie) was prepared at a concentration of 1.5×10^5^ cells/ml. 150 µl of cell suspension was mixed with 150 µl of basal medium containing DMSO (control), recombinant human VEGF-C (hVEGF-C, 100ng/ml) (Cat# RLT-300-079, Axxora, San Diego, CA), hTGF-β1 (1ng/ml)(Cat# 240-B, Bio-Techne/ R&D Systems), and TGF-β inhibitor (3uM) (LY364947, Cat#L6293, Sigma-Aldrich) as indicated in Figure 5G&H, and loaded on each Matrigel® coated well. The plate was incubated in a humidified incubator for 4 hours at 37°C. Five DIC images at x100 magnification were obtained per well using Zeiss Axio Observer. The total length of cords was measured by ImageJ (NIH) using Angiogenesis Analyzer macro (Gilles Carpentier in 2012; http://image.bio.methods.free.fr/ImageJ/?Angiogenesis-Analyzer-for-ImageJ&lang=en).

### BrdU Cell Proliferation Assay

Cell proliferation was assessed by a BrdU cell proliferation ELISA kit (Cat#ab126556, Abcam) according to the manufacturer’s instruction. Briefly, Human liver LyECs were plated in 96-well plate overnight and then treated with DMSO (control), hVEGF-C (100ng/ml) (Cat# RLT-300-079, Axxora), hTGF-β1 (1 ng/ml) (Cat# 240-B, Bio-Techne/ R&D Systems), TGF-β inhibitor (3uM) (LY364947, Cat#L6293, Sigma-Aldrich) alone or together in BrdU-containing ENDO-Growth medium (Cat# cAP-05, Angio-Proteomie) for 6 hours (Figure 5I). After treatment, cells were fixed and incubated with anti-BrdU antibody for 1 hour at room temperature. After incubation, cells were washed 3 times in the wash buffer and then incubated with peroxidase goat anti-mouse IgG conjugate for 30 minutes at room temperature and washed for 3 times. Finally, TMB Peroxidase Substrate was added to cells and incubate for 30 minutes at room temperature in dark before adding the Stop Solution. The plate was read at dual wavelengths of 450/550 nm, and proliferation rates were calculated by comparing the increased rate with the untreated cells.

### Quantitative real-time polymerase chain reaction

Total RNA was extracted by TRIzol reagent (Invitrogen) from primary human liver LyECs according to the manufacturer’s instruction. RNA quantity and quality were assessed by micro-volume spectrophotometry (NanoDrop 2000, Thermo Fisher Scientific). Total RNA (700 ng) was reverse transcribed into cDNA using a Reverse Transcription Reagents kit (Cat#04897030001, Roche Diagnostics, Indianapolis, IN). Quantitative real-time PCR (qPCR) was performed for cDNA samples using iTaq Universal SYBR Green (Cat#P1725121, Bio-Rad Laboratories) on the ABI 7500 Real-Time PCR System (Applied Biosystems) or the QuantStudio 6 Flex Real-Time PCR System (Applied Biosystems). 18S was used as a loading control. Results are shown as fold-changes relative to control groups. Primer sequences are listed below: 18S (forward: GCAATTATTCCCCATGAACG, reverse: GGCCTCACTAAACCATCCAA), Mouse VEGF-C (forward: TGTCTCTGGCGTGTTCCCT, reverse: ATCAGCTCATCTACGCTGGAC), and Human PAI-1 (forward: AGAGGTGGAGAGAGCCAGAT, reverse: TCCACTGGCCGTTGAAGTAG)

### Western blot analysis

Western blots were performed as previously described(24). Primary antibodies, including [Collagen I (COl I, dilution factor 1:1000, Cat# ab292, Abcam, Cambridge, MA), Collagen III (COl III, 1:1000, Cat#NBP2-15946, NOVUS Biological), Fibronectin I (FN1, 1:1000, Cat#ab2413, Abcam), aSMA (1:2000, Cat#ab124964, Abcam), GAPDH (1:1000, Cat#2118, Cell Signaling Technology), pSMAD2/3 (1:1000, Cat# 8828, Cell Signaling Technology), Total SMAD2/3 (1:1000, Cat#3102, Cell Signaling Technology)], were applied on 0.2 μm nitrocellulose membranes (Cat#1620112, Bio-Rad Laboratories, Hercules, CA). Fluorophore-conjugated secondary antibodies (Li-Cor Biotechnology, Lincoln, NE) having 680 or 800 nm emission were used for detection. Protein bands were visualized and quantified using the Odyssey Infrared Imaging System (Li-Cor Biotechnology).

### Mouse VEGF-C ELISA

Mouse VEGF-C level was determined in homogenized mouse liver tissue using a mouse VEGF-C ELISA Kit (Cat# OKEH00240, Aviva Systems Biology, San Diego, CA). All procedures were followed according to the manufacturer’s instructions.

### Statistical analysis

All data are expressed as mean ± standard error of mean (SEM). The comparison between two groups was performed using unpaired Student’s t-test. The comparison among multiple groups was performed using one-way ANOVA. A *p*-value less than 0.05 was considered statistically significant. Kaplan–Meier analysis followed by a log-rank test with or without post hoc Bonferroni’s correction was used to compare survival between mouse groups.

## Data availability statement

The raw data that support the findings of this study have been assigned GEO accession numbers (GSE276093). The GSE records are scheduled to be publicly available on Dec 31, 2024.

## Supplementary figures

**Supplementary Figure 1.**
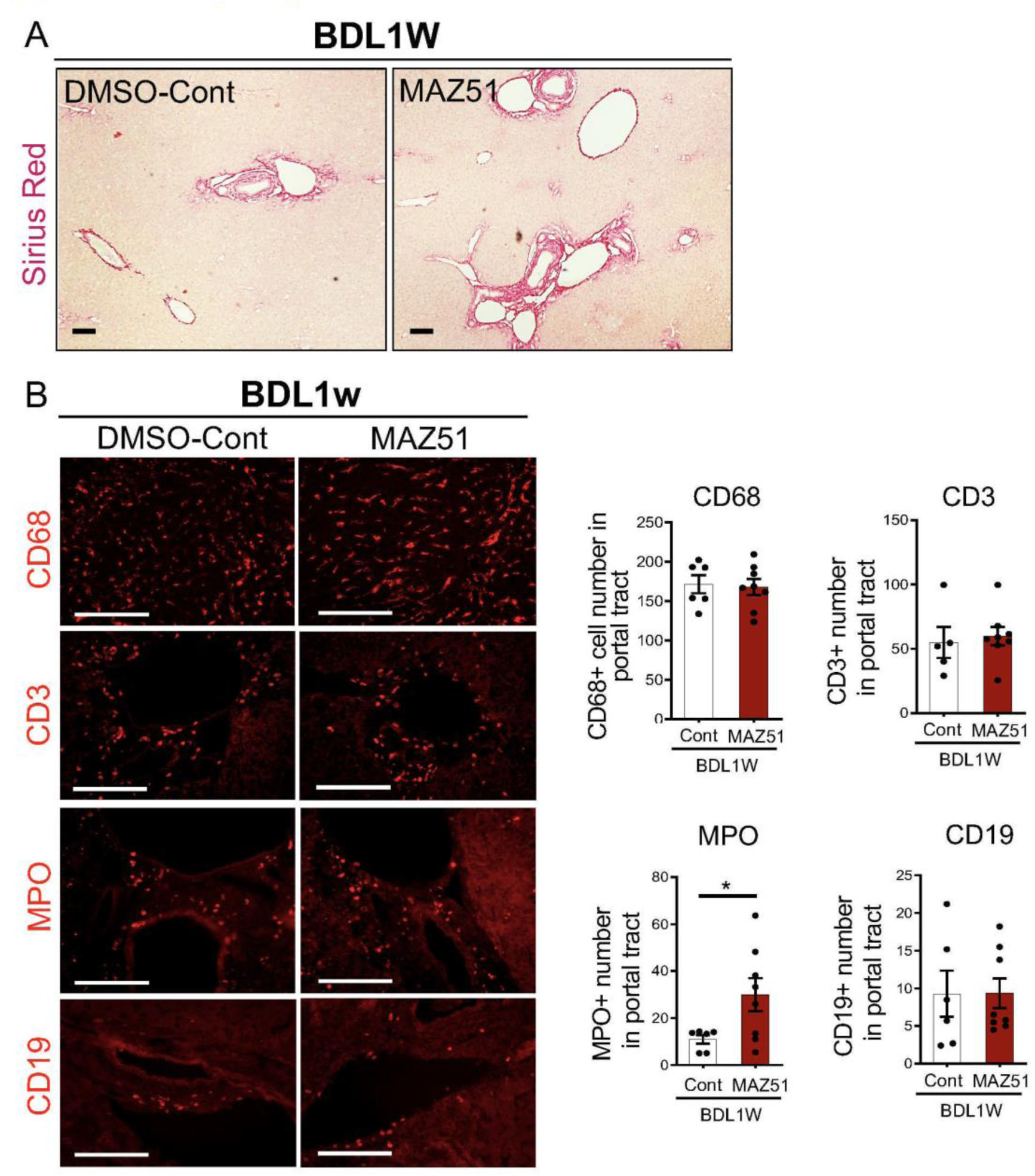
Fibrosis and immune cell Infiltration in livers from BDL1w mice treated with DMSO (Control) or MAZ51. (A) Representative Sirius red staining (larger view) images. (B) Immunofluorescence (IF) images and quantification of CD68 (macrophage; red), CD3 (T cell; red), MPO (neutrophil; red) and CD19 (B cell; red) in liver from control (n=7) and MAZ51 (n=7) treated 1-week BDL mice. *p < 0.05. Scale bars: 100 μm.

**Supplementary Figure 2.**
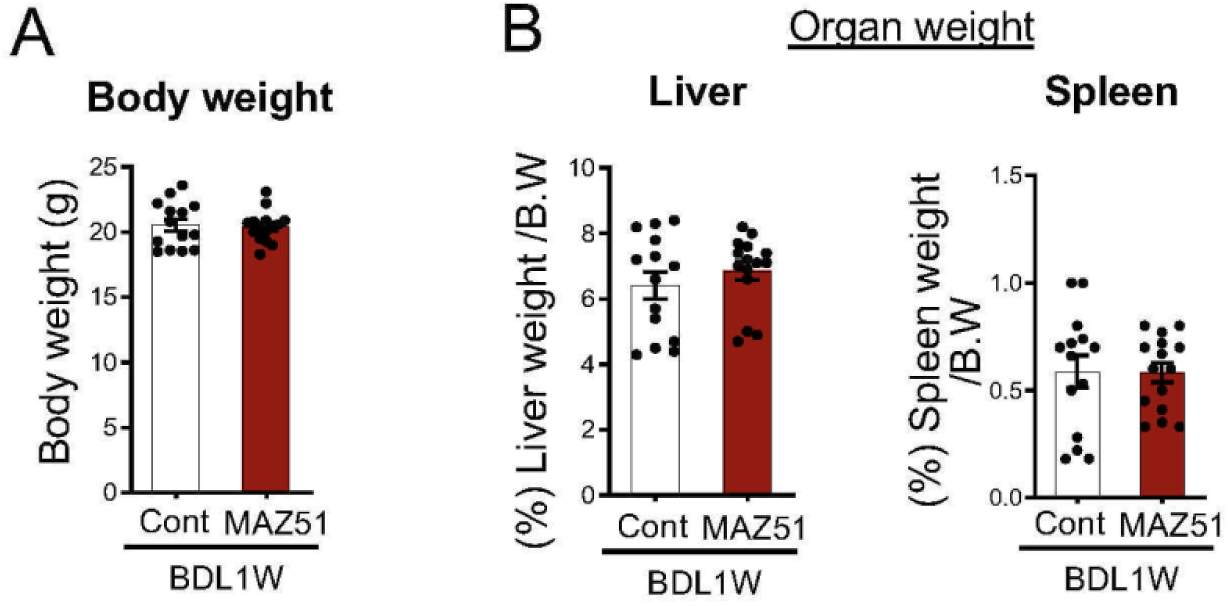
Body weight and percent organs weight ratio of BDL1w mice treated with DMSO (Control) or MAZ51. (A) Body weight (B.W.), (B) liver weight and spleen weight in mice treated with DMSO (Control, n=11) or MAZ51 (n=9).

**Supplementary Figure 3.**
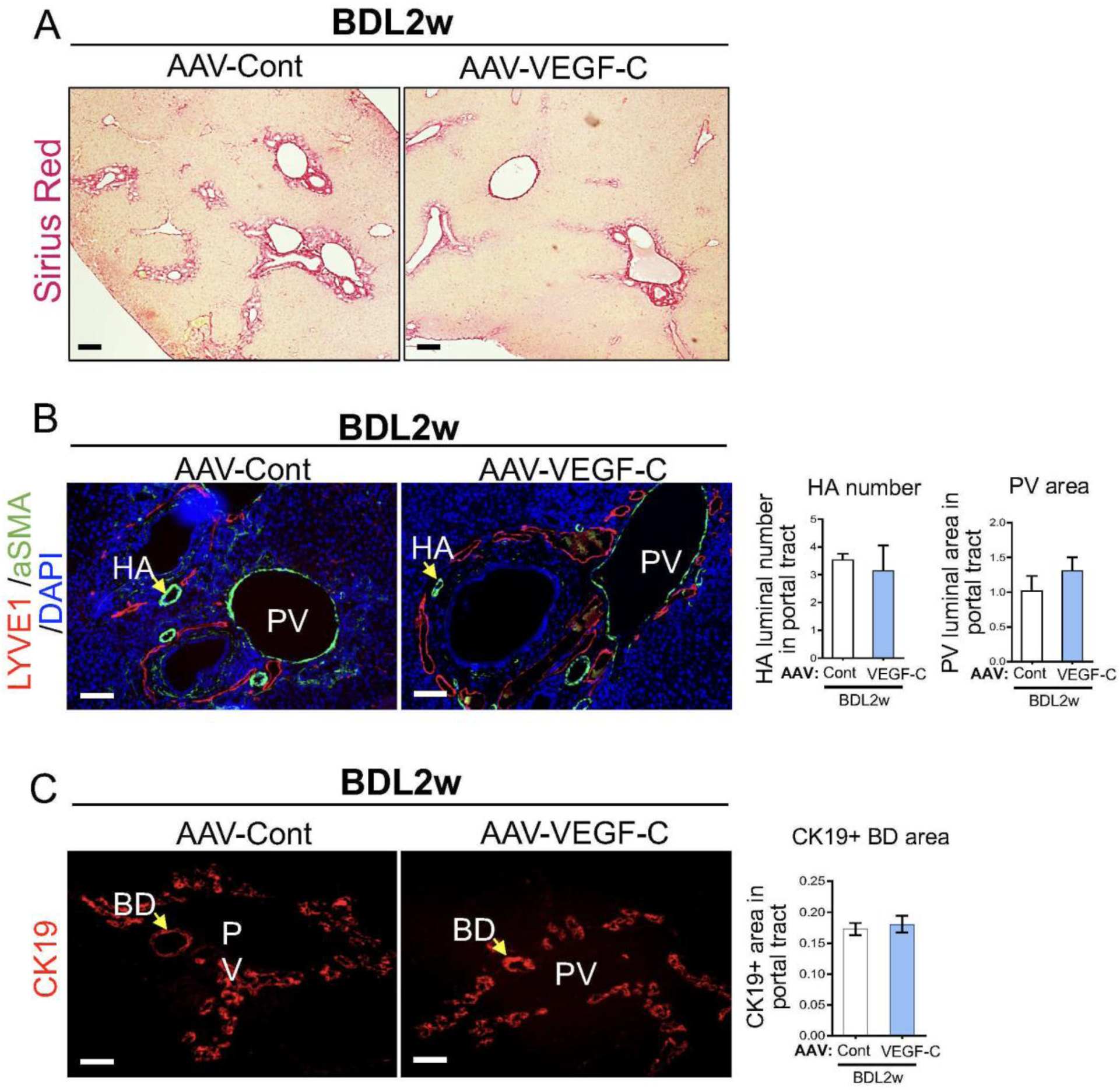
VEGF-C overexpression reduces liver fibrosis without inducing blood vessel and bile duct expansions in the liver from BDL2w mice treated with AAV8-VEGF-C. (A) Representative images of Sirius red staining. (B) Immunofluorescence (IF) images and quantification of hepatic artery (HA) number and portal vein (PV) area in livers from control(n=7) and AAV8-VEGF-C (n=7) treated 2-week BDL mice. (C) IF images and quantification of bile duct (CK19; red) in livers from control(n=7) and AAV8-VEGF-C(n=7)-injected 2-week BDL mice. Scale bars: 100μm.

**Supplementary Figure 4.**
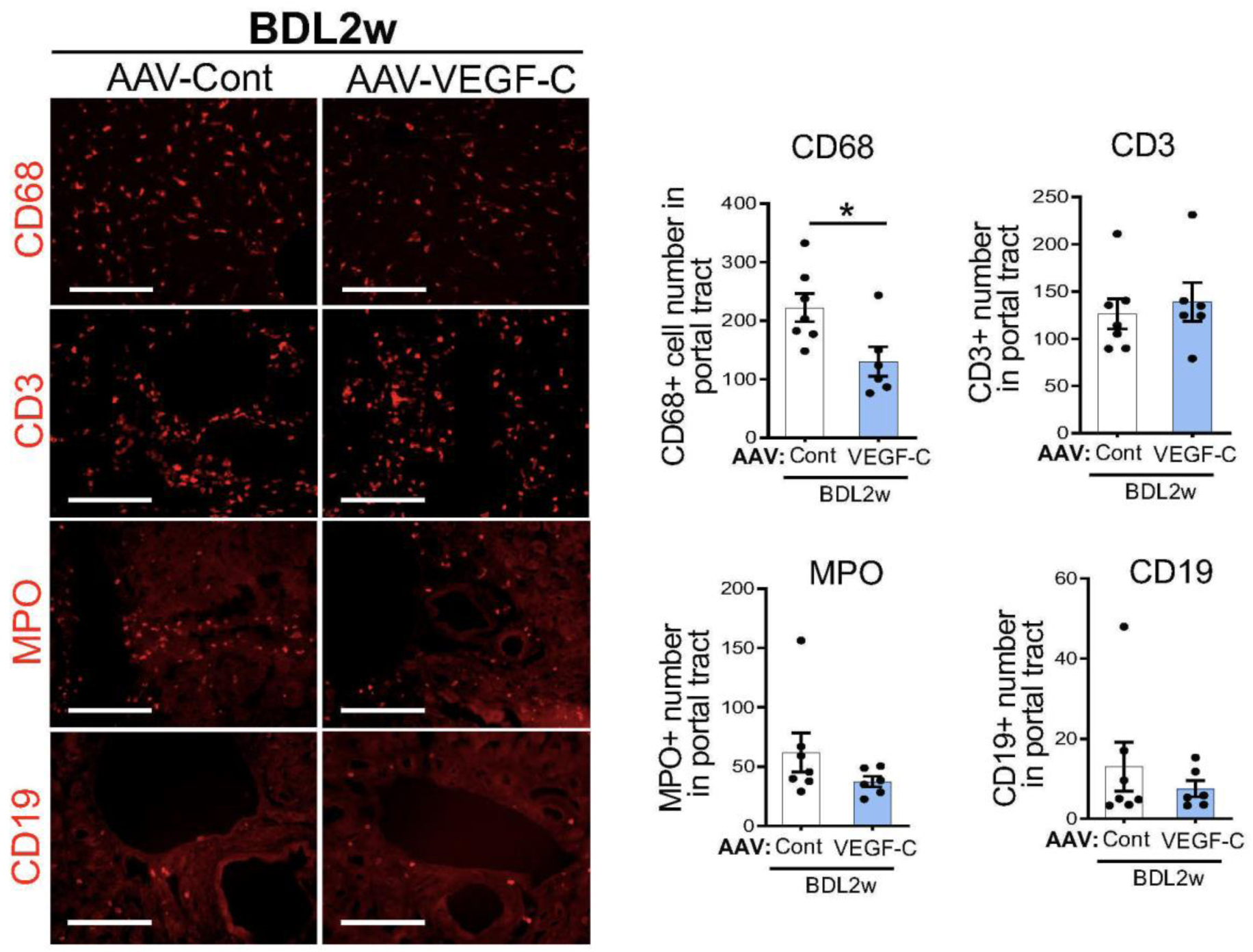
Immune cell infiltration in livers from BDL2w mice treated with AAV8-GFP (Control) or AAV8-VEGF-C. Immunofluorescence (IF) images and quantification of CD68 (macrophages; red), CD3 (T-cells; red), MPO (neutrophils; red) and CD19 (B cells; red) in livers from control (n=7) and AAV8-VEGF-C (n=7) treated 2-week BDL mice. *p< 0.05. Scale bars: 100 μm.

**Supplementary Figure 5.**
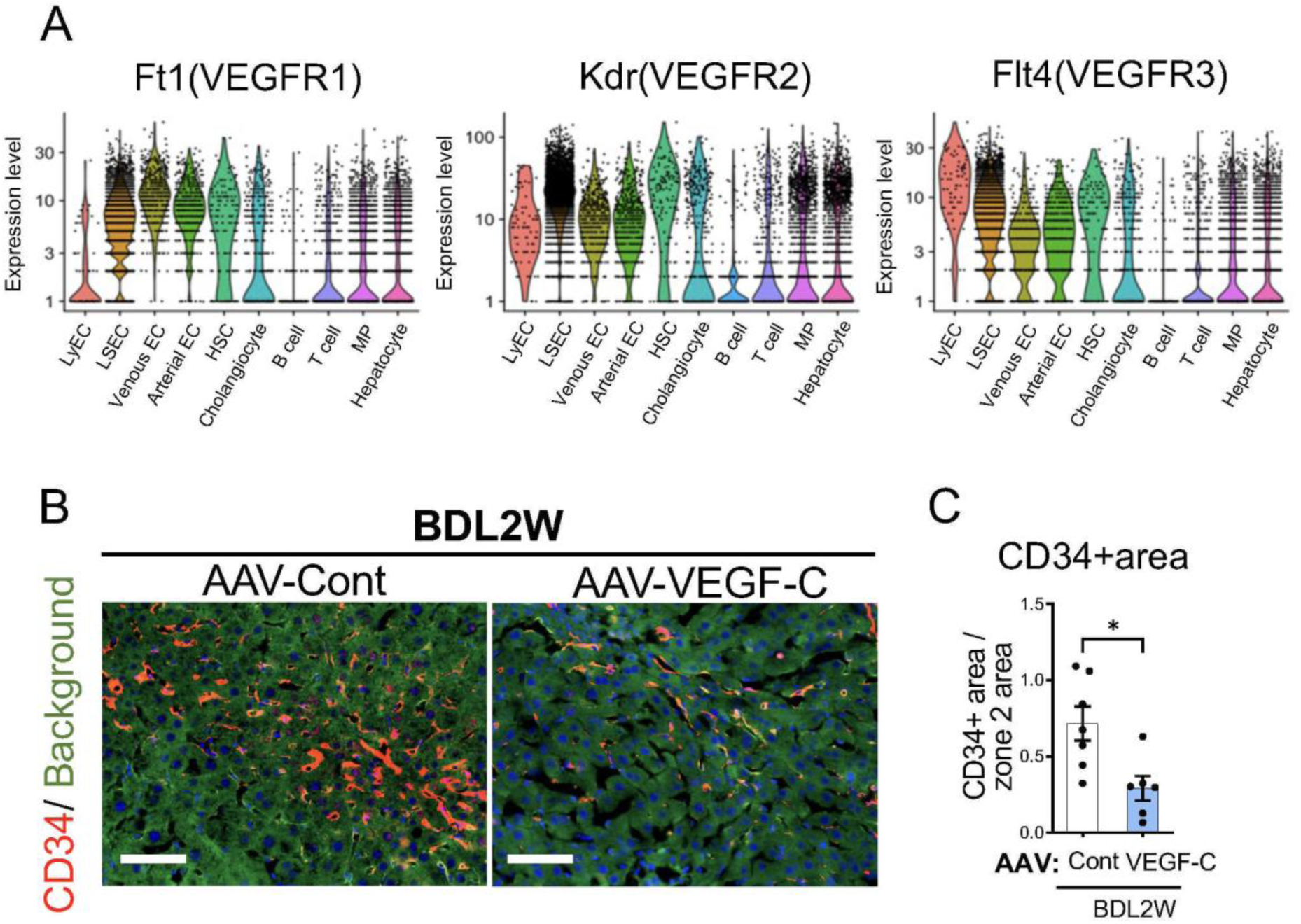
VEGF-C overexpression ameliorates capillarization of liver sinusoidal endothelial cell (LSEC) in cholestatic mice. (A) Violin plots of gene expression of *Flt1*, *Kdr*, and *Flt4* in livers from healthy(n=3) and cirrhotic(n=3) mice (Cdh5-Cre-mTmG^+/+^ mice) (GSE147581). (B) Immunofluorescence (IF) images and (C) quantification of CD34 (a marker of LSEC capitalization; red) in livers from control(n=7) and AAV8-VEGF-C (n=7) administered 2-week BDL mice. *p< 0.05.

**Supplementary Figure 6.**
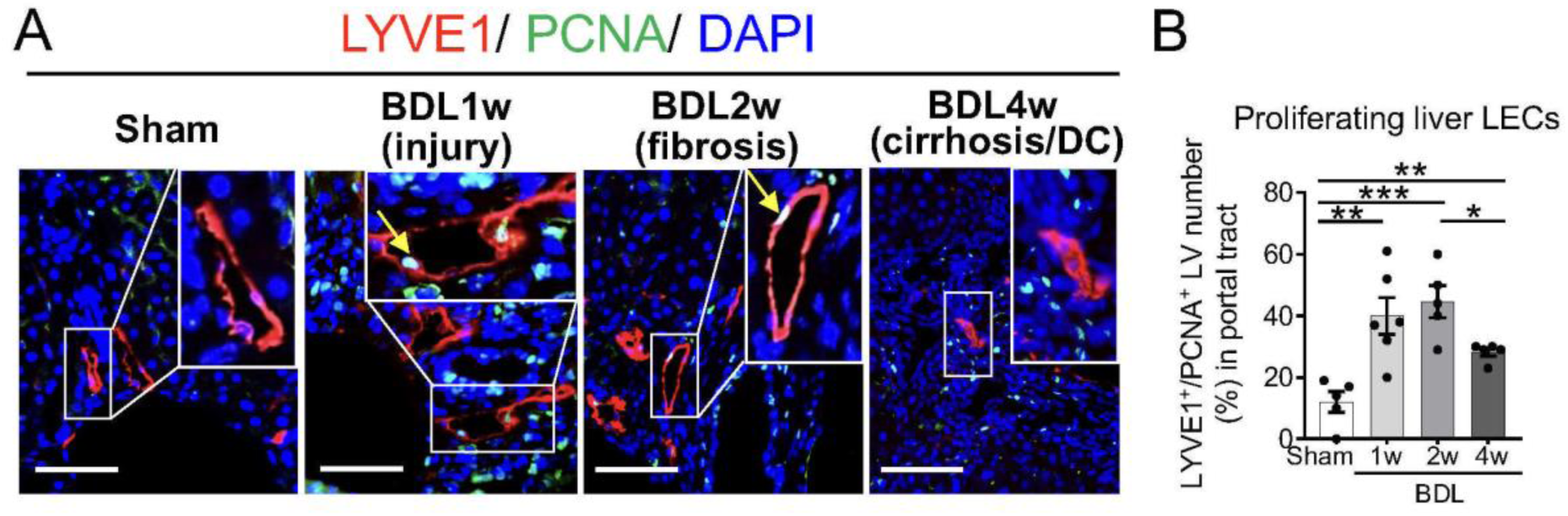
Proliferating liver lymphatic endothelial cells (LyECs) are increased in the early stage of fibrosis/cirrhosis but decreased in the late-stage cirrhotic BDL mice. (A) Representative immunofluorescence (IF) images of proliferating cell nuclear antigen (PCNA; green), LYVE1 (red) and DAPI in sham, 1-week, 2-week, 4-week BDL mouse livers. PCNA-positive nuclei with DAPI in LYVE 1 positive LVs (arrows) are proliferating lymphatic endothelial cells (LyECs). (B) Quantification of PCNA-positive LyECs in livers from sham(n=5), 1-week(n=5), 2-week(n=5), and 4-week(n=7) BDL mice. *p< 0.05, **p< 0.01, and ***p< 0.001. Scale bars: 100 μm. PV, portal vein.

**Supplementary Figure 7.**
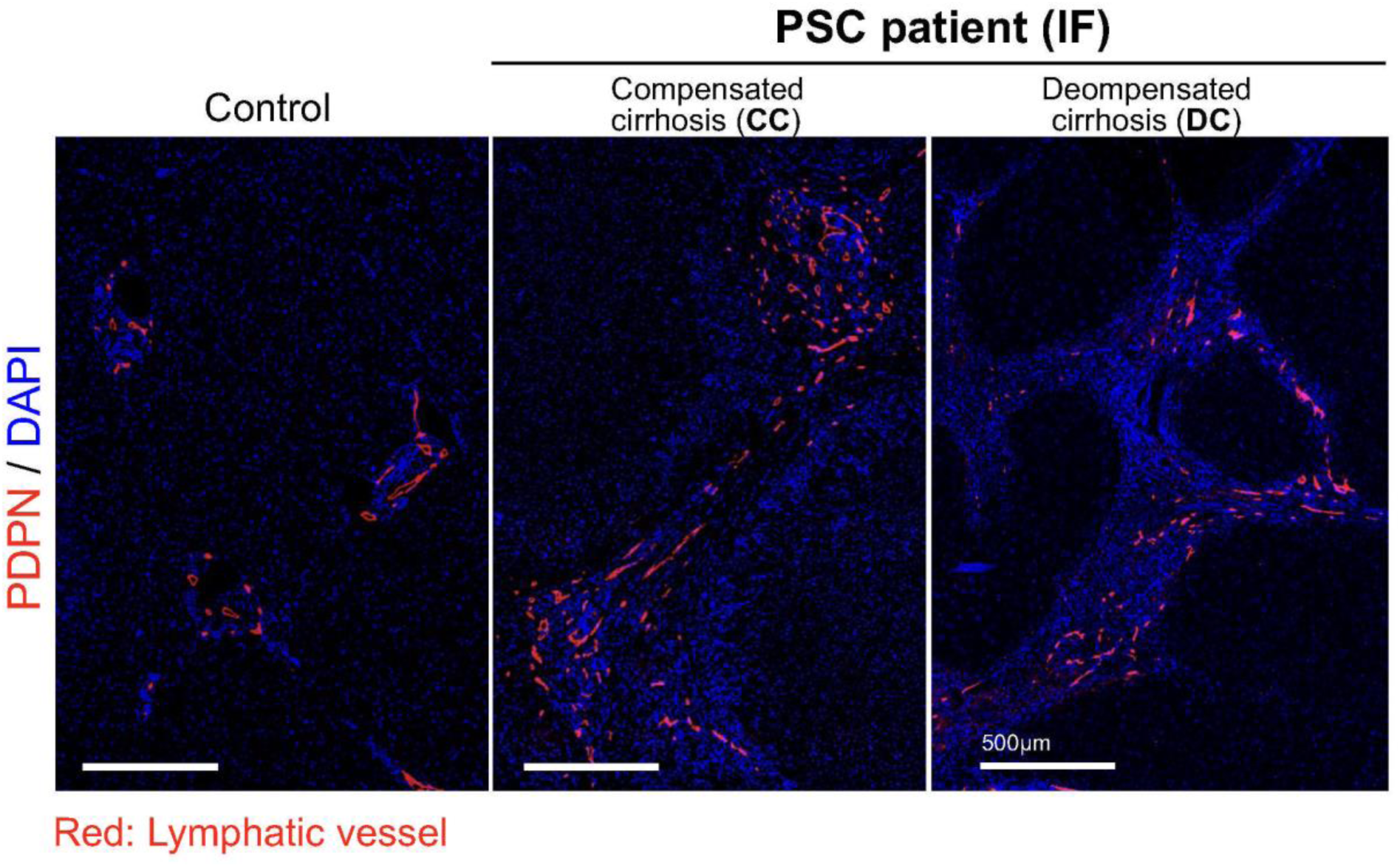
Liver lymphatic vessel (LV) numbers and area are increased in compensated cirrhosis (CC) but decreased at decompensated cirrhosis (DC) in primary sclerosing cholangitis (PSC) patients. Tile scan images of podoplanin (PDPN, a marker of LVs; red) immunofluorescence (IF) staining in livers specimens from healthy controls liver and patients with primary sclerosing cholangitis (PSC) including CC and DC livers.

**Supplementary Figure 8.**
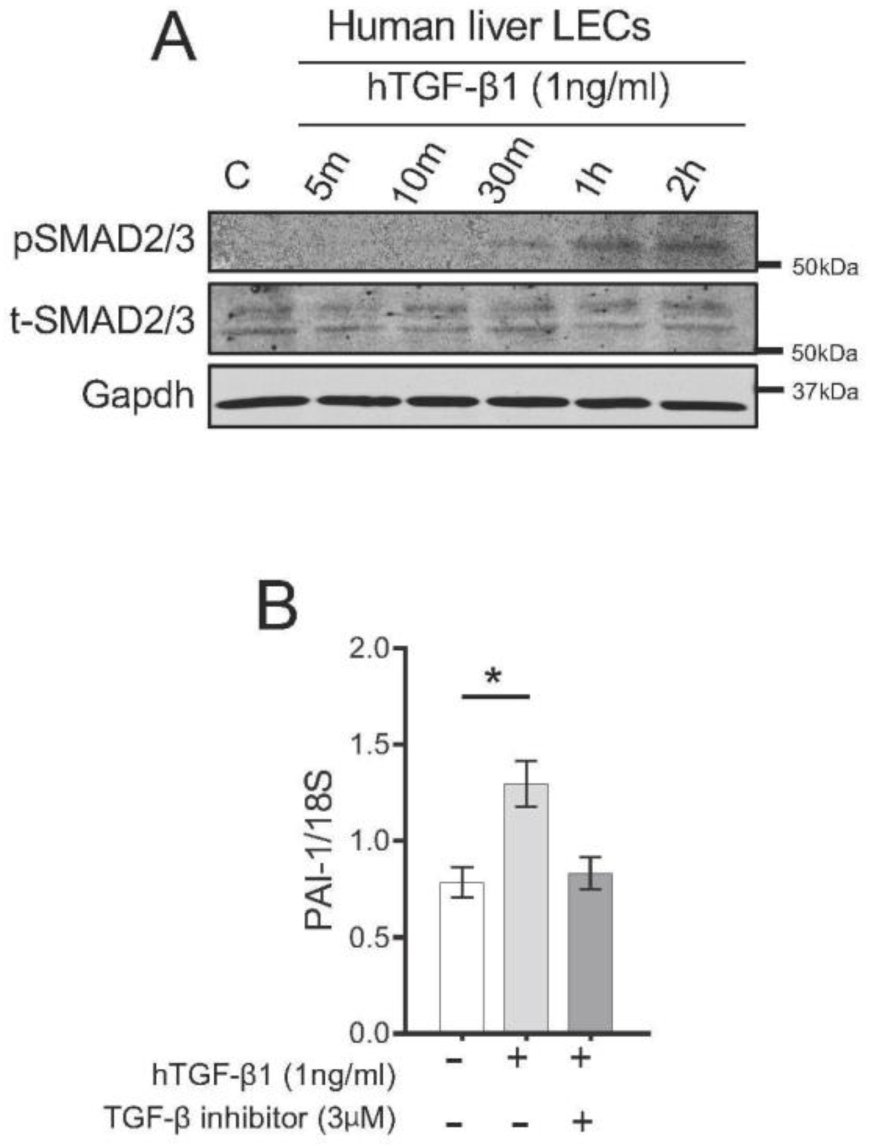
TGFβ signaling transduction in human liver lymphatic endothelial cells (LyECs) **(A)** Phospho-SMAD2/3 levels in human primary liver LyECs treated with human TGF-β1(hTGFβ1, 1ng/ml) at indicated time. **(B)** Quantification of PAI-1 mRNA expression in human primary liver LyECs treated with human recombinant TGF1 or control-DMSO for 24 hours. Representative data from three independent experiments. *p< 0.05.

**Supplementary Figure 9.**
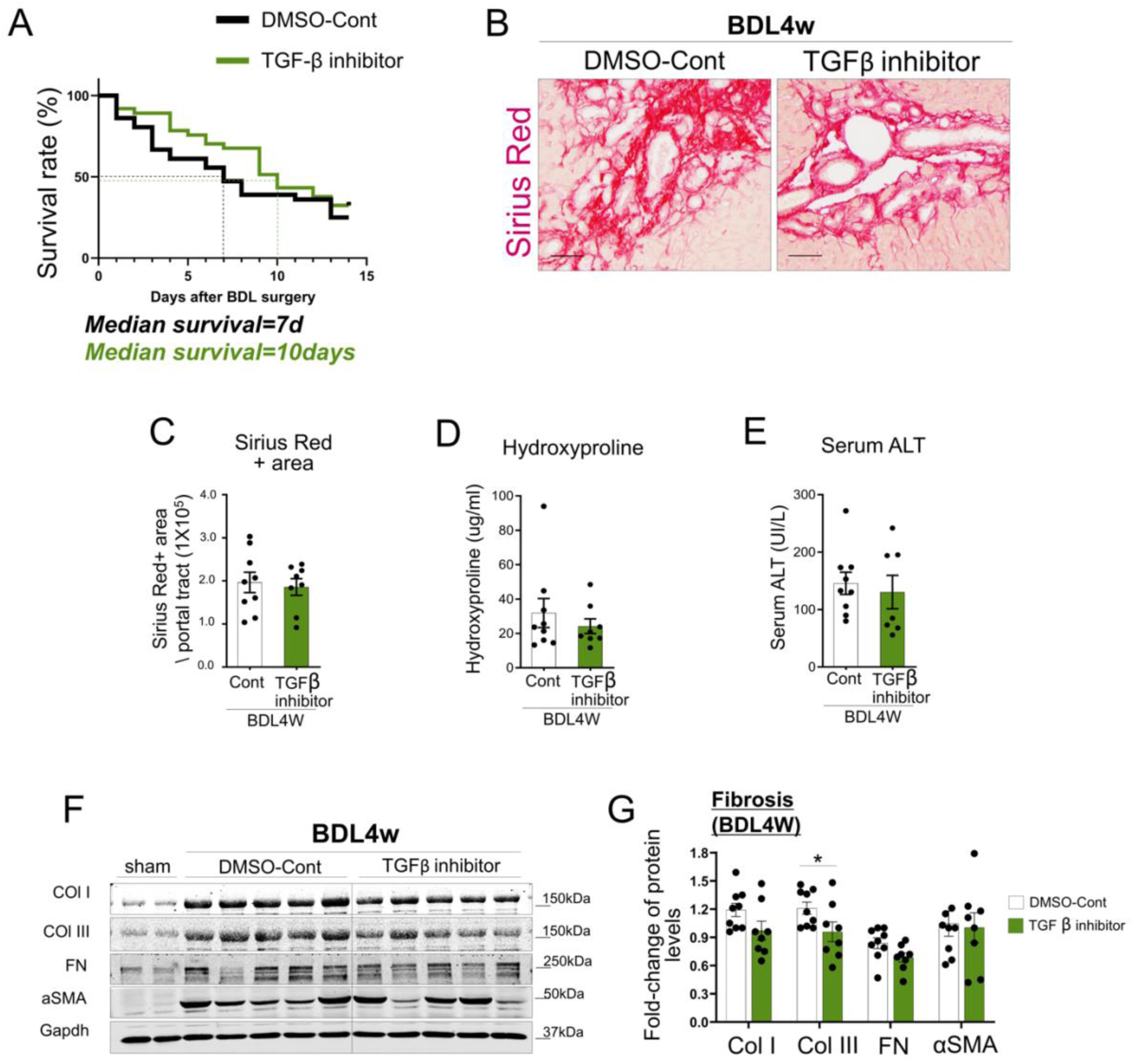
Administration of TGF-β inhibitor prolonged median survival rate in late-stage cirrhotic mice. (A) Kaplan-Meier survival curve showing a statistically significant difference in median survival as indicated from 4-weeks BDL mice treated with control-DMSO(n=36) or TGF-β inhibitor(LY364947, 0.02mg/mouse, n=37). (B-G) Evaluation of liver fibrosis and injury. Representative Sirius red staining images (B) and its quantification (C), as well as hydroxyproline measurement (D) and serum ALT levels (E) in livers from 4-week BDL mice treated control-DMSO(n=8) and TGF-β inhibitor(LY364947, n=7). (F& G) Levels of liver fibrosis makers including collagen I (ColI), collagen III (Col III), fibronectin 1 (FN1) and α-SMA. Control-DMSO treated (n=8) or TGF-β inhibitor treated (n=7) 4-week BDL mice. *p< 0.05. Scale bars: 100 μm. PV, portal vein.

